# Branched chain amino acid synthesis is coupled to TOR activation early in the cell cycle in yeast

**DOI:** 10.1101/2023.01.10.523468

**Authors:** Heidi M. Blank, Carsten Reuse, Kerstin Schmidt-Hohagen, Staci E. Hammer, Karsten Hiller, Michael Polymenis

## Abstract

How cells coordinate their metabolism with division determines the rate of cell proliferation. Dynamic patterns of metabolite synthesis during the cell cycle are unexplored. We report the first isotope tracing analysis in synchronous, growing budding yeast cells. Synthesis of leucine, a branched-chain amino acid (BCAA), increased through the G1 phase of the cell cycle, peaking later during DNA replication. Cells lacking Bat1, a mitochondrial aminotransferase that synthesizes BCAAs, grew slower, were smaller, and were delayed in the G1 phase, phenocopying cells in which the growth-promoting kinase complex TORC1 was moderately inhibited. Loss of Bat1 lowered the levels of BCAAs and reduced TORC1 activity. Exogenous provision of BCAAs to cells lacking Bat1 promoted cell division and increased TORC1 activity. In wild-type cells, TORC1 activity was dynamic in the cell cycle, starting low in early G1 but increasing later in the cell cycle. These results suggest a link between BCAA synthesis from glucose to TORC1 activation in the G1 phase of the cell cycle.

## INTRODUCTION

How metabolism is coupled to cell division underpins the control of cell proliferation. Except for early embryonic cell divisions, the rate at which cells can divide usually depends not on how fast they can duplicate their genome but on how fast they can synthesize everything else that makes up a newborn cell. Since the discovery of *cdc* mutants in *S. cerevisiae* decades ago, it has been known that cells continue to grow if cell division is blocked (Johnston *et al*, 1977; Hartwell & Unger, 1977). On the other hand, stopping cell growth blocked cell division. In some instances, the cell cycle machinery does trigger specific metabolic pathways, especially in lipid and carbohydrate mobilization (Ewald *et al*, 2016; Kurat *et al*, 2009). By and large, however, metabolism controls division and not the other way around. Similarly, inhibiting the target of rapamycin (TOR) kinase, the ‘master regulator’ of multiple anabolic pathways, including protein synthesis, arrests cells in early G1 with a small size (Barbet *et al*, 1996).

Early reports of cell cycle-dependent metabolic changes in a variety of organisms were later shown to be artifacts of synchronization (Creanor & Mitchison, 1979; Creanor *et al*, 1983). Popular methods to synchronize cells rely on first blocking them at some point in the cell cycle and, after some time, releasing the block so cells can progress synchronously in the subsequent phases of the cell cycle. But cells continue to grow during arrest, perturbing the physiological coupling between cell growth and division. In recent years, several studies monitored the steady-state levels of metabolites in the cell cycle in yeast (Ewald *et al*, 2016; Blank *et al*, 2020), green alga (Jüppner *et al*, 2017), fly (Sanchez-Alvarez *et al*, 2015), and human cells (Atilla-Gokcumen *et al*, 2014; Scaglia *et al*, 2014).

However, metabolite abundances do not reveal material flow per unit time through a metabolic pathway (Jang *et al*, 2018). One can infer that information through the use of isotope tracers and the appropriate computational analyses (Weindl *et al*, 2015). Isotope tracing in the cell cycle to monitor the flow through metabolic pathways has remained unexplored. Only one study reported Kreb’s cycle flux in dividing human HeLa cells (Ahn *et al*, 2017), synchronized with a double thymidine block, which arrests cells at the G1/S transition (Polymenis, 2022). When the cells were released from their arrest, glucose oxidation peaked late in the subsequent G1 phase, followed by oxidative and reductive glutamine metabolic fluxes in S phase (Ahn *et al*, 2017). It is unknown whether these metabolic changes extend to other systems and how they are connected to molecular pathways that govern cell growth and division.

Here, we queried for the first time with isotope tracing highly synchronous, unarrested, dividing yeast cells. We identified an increase in the synthesis of the branched-chain amino acid (BCAA) leucine in the G1 phase of the cell cycle, and peaking later in the S phase. We found that cells lacking the mitochondrial BCAA aminotransferase Bat1 were smaller, had a much longer G1 phase, and very low TOR complex 1 (TORC1) activity. In newborn wild-type daughter cells, TORC1 activity was also low in early G1 but increased as cells progressed in the cell cycle. These results provide the first picture of cell cycle-dependent metabolic fluxes in yeast, potentially linking BCAA synthesis to TORC1 activation in the cell cycle.

## RESULTS AND DISCUSSION

### Experimental design of isotope tracing in the cell cycle

We used a prototrophic strain, CEN.PK (see Materials and Methods), cultured in synthetic minimal medium (SMM) to avoid biases that could arise from exogenous nutrient supplementation. To minimize arrest-related artifacts, we used centrifugal elutriation to isolate unarrested, small, early G1 cells (Polymenis, 2022). Using uniformly labeled ^13^C-glucose we generated a sliding pulse-chase window over the entire cell cycle (see Figure 1A and Materials and Methods).

**FIGURE 1.**
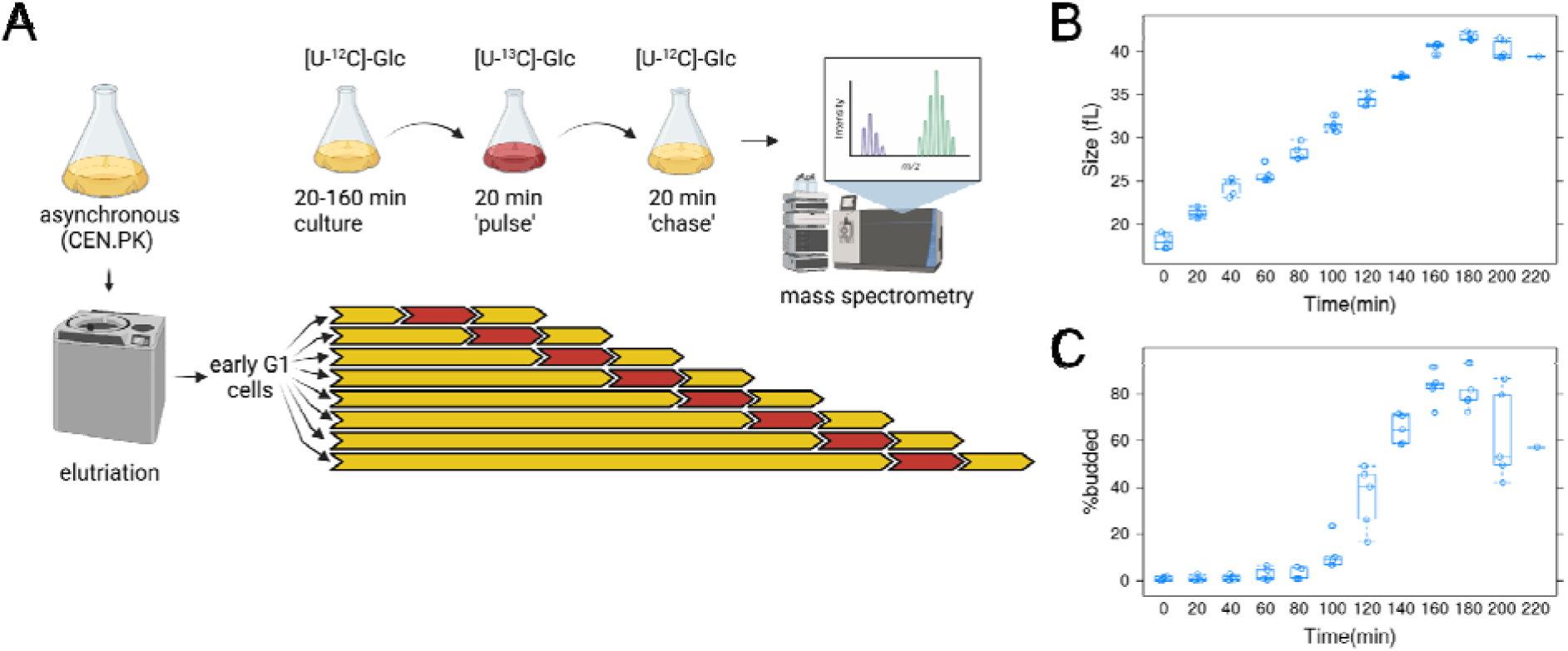
Overview of the approach to obtain samples for metabolic flux analysis in highly synchronous, dividing, budding yeast cells. A, For each experiment, early G1 daughter cells of a prototrophic strain (CEN.PK; see Materials and Methods) were obtained by centrifugal elutriation. The elutriated culture was split into eight aliquots and cultured for a varying amount of time, from 20 to 160 min, in minimal, [U-^12^C]-glucose medium. They were then transferred to a medium with [U-^13^C]-glucose for 20 min (pulse) and then incubated for another 20 min in [U-^12^C]-glucose medium (chase). Metabolite extracts from these cells were analyzed by mass spectrometry. Five such independent experiments were performed. The figure was generated with Biorender.com. B, Boxplots showing the cell size (y-axis) over time (x-axis) of all the samples as they progressed in the cell cycle. C, Boxplots showing the percentage of budded cells (y-axis) from the same samples shown in B. The boxplot graphs were generated with R language functions. The values used to draw the plots are in File S1/Sheet1.

To gauge cell cycle progression and synchrony, we measured cell size with a channelyzer (Figure 1B) and the percentage of budded cells with microscopy (Figure 1C). In *S. cerevisiae* the appearance of a bud on the cell surface correlates with the timing of initiation of DNA replication (Hartwell & Unger, 1977; Pringle, 1981). Cells progressed synchronously in the cell cycle, increasing in size at a constant rate (Figure 1B), with buds appearing at ∼100 min, while by 160-180 min the vast majority of cells were budded and had doubled in size (to >40fL; Figure 1C). By 180 min, new G1 daughter cells were generated, leading to a drop in the percentage of budded cells and cell size (Figure 1B,C). These data suggest that the metabolically labeled samples we prepared were from highly synchronous cells, querying all cell cycle phases.

### Synthesis of leucine from glucose changes in the cell cycle

Analyzing the metabolically labeled cell extract samples by gas chromatography-mass spectrometry (GC-MS) revealed changes in the mass isotopomer distributions (MIDs) in some of the species we detected (Figure 2 and File S2). MID changes reflect patterns in the synthesis and turnover of the relevant metabolites. Trends of dynamic changes in isotopomer abundance were evident for several species (e.g., pyruvate (M2), succinate (M3), Figure 2). However, based on ANOVA statistical analysis, leucine (M6) and pyruvate (M1) were the most significantly affected metabolites we detected (Figure 2 and FileS2/Sheet2). Pyruvate (M1) likely arises from breakdown pathways. For the synthesis of leucine, two molecules of ^13^C-labeled pyruvate and one molecule of ^13^C-labeled acetyl-CoA provide the carbon input from the glycolytic breakdown of the ^13^C-labeled glucose to the leucine (M6) isotopomer (see Figure EV1). The levels of leucine (M6) were the lowest at the earliest time point in early G1 but increased (>4-fold) by 100 min, peaking at ∼150 min in the S phase (Figure 2), when more than half the cells in that experiment were budded (Figure 1C). These results are consistent with an increase in newly synthesized leucine from glucose as cells progress through the G1 phase of the cell cycle.

**FIGURE 2.**
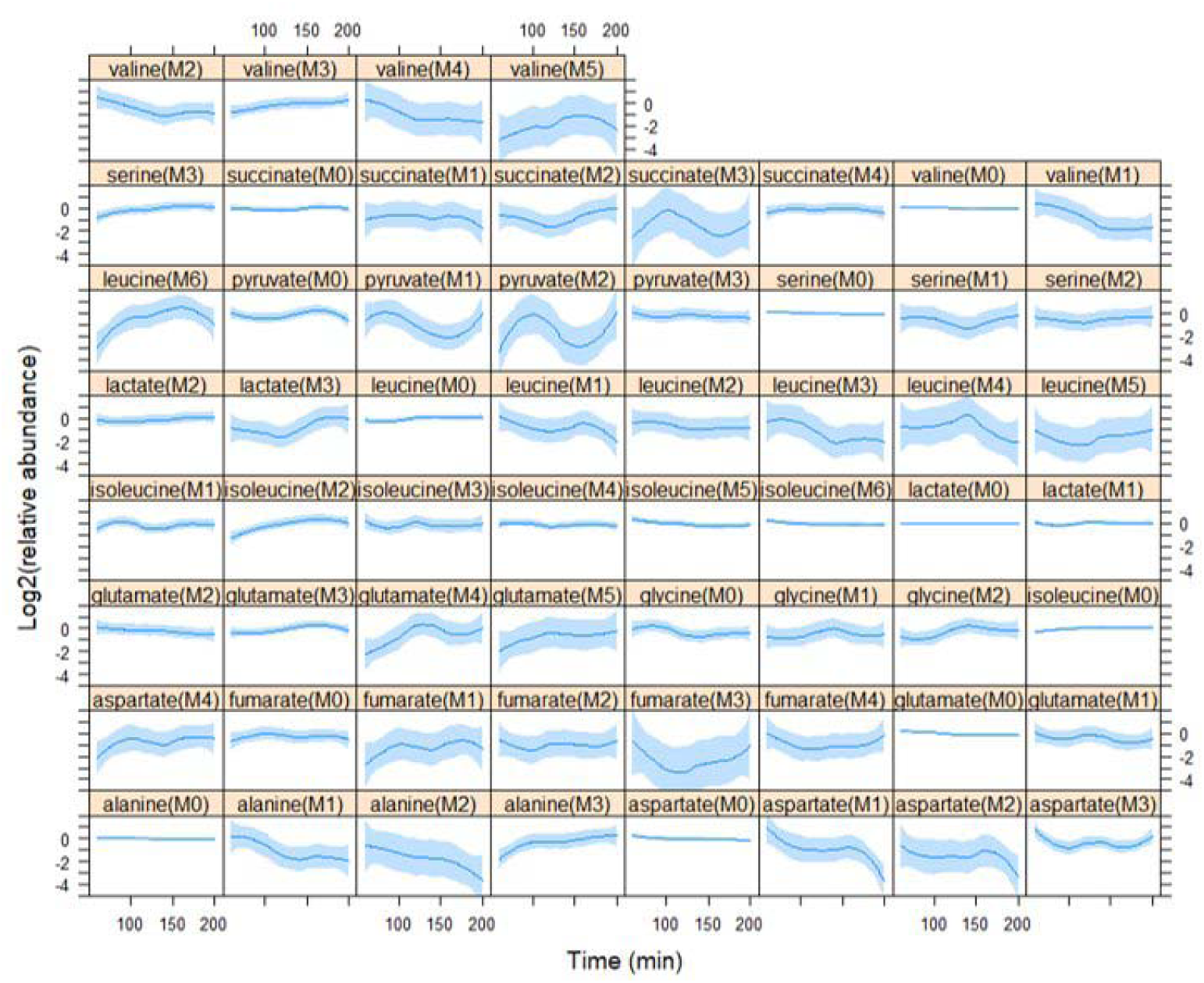
Relative abundance of intracellular metabolite isotopomers in the cell cycle. Each plot shows the relative mass isotopomer distribution (MID) values for metabolite species detected from cell extracts in each of the five independent experiments shown in Figure 1. Each value was divided by the average value of the entire time series (x-axis) for that species from the same experiment, and Log2-transformed (y-axis). Loess curves and the std errors at a 0.95 level are shown. The values used to generate the graphs are in File S1/Sheet2. The Log2(relative abundance) values of pyruvate(M1) and leucine(M6) showed the most significant changes (p<0.1, see File S2/Sheet2) in cells across the cell cycle.

We also measured the MID values from the media of the same cultures used to prepare the cell extracts (File S2/Sheet3). The levels of the glutamate (M4) and glutamine (M2) forms in the media were significantly changed in the cell cycle (File S2/Sheet4). Still, these changes were due to a single time point of the experiment (e.g., glutamate (M4) levels in the media were the highest at the first time point we measured, compared to all subsequent time points), making interpretations difficult. All the MID measurements and data collected on selected ion monitoring (SIM) mode are in File S2.

Whether changes in the synthesis of metabolites identified from isotope tracing experiments are reflected in their steady-state levels depends on several variables. These variables include the metabolite turnover, partition in different pools inside cells, and their generation from the breakdown of macromolecules (e.g., from proteins, in the case of amino acids). We measured the intracellular levels of amino acids under the same experimental conditions as in Figure 1. Based on a phenylthiohydantoin (PTH)-based detection method (see Materials and Methods), the steady-state levels of several amino acids changed in the cell cycle, but in no case varying >2-fold (Figure EV2). For example, arginine and asparagine, which were not detected in our isotope tracing experiment, peaked at different times in the cell cycle, in G2/M vs. G1, respectively (Figure EV2). The levels of the BCAA amino acids (Ile, Leu, Val) start high but decline in late G1, when the cells reach ∼35 fL, then increase by ∼30-40% in G2/M (Figure EV2). Although the same trend was evident for all BCAAs, these differences were significant only for valine (p<0.05, based on robust ANOVA analysis) because the variance in the measurements of leucine and isoleucine was higher.

In the remainder of this manuscript, we focus on the role of BCAA synthesis in proliferating cells. We note that although leucine (M6) levels changed the most significantly in the cell cycle, valine (M5) levels also showed a similar trend, peaking when cells were in the S phase (see Figure 2, top right panel).

### Cells lacking Bat1 grow slowly

The BCAA aminotransferases Bat1 and Bat2 catalyze the synthesis of all BCAAs from the corresponding alpha-keto acids (Figure 3A) (Kispal *et al*, 1996; Eden *et al*, 1996). Bat1 is mitochondrial, while Bat2 is cytoplasmic (Kispal *et al*, 1996; Eden *et al*, 1996; Prohl *et al*, 2000). We deleted *BAT1* or *BAT2* in the same prototrophic strain background (CEN.PK; see Materials and Methods) we used in the isotope tracing experiments. There are conflicting reports in the literature about the growth rate of *bat1*Δ cells. One of the studies that identified *BAT1* reported that *bat1*Δ cells grew and divided faster (Schuldiner *et al*, 1996). However, other studies reported that loss of Bat1 reduced growth rate, especially in minimal media (Kispal *et al*, 1996; Takpho *et al*, 2018; Koonthongkaew *et al*, 2020). We found that cells lacking Bat1, but not Bat2, grew poorly on minimal media (Figure 3B). Exogenous supplementation of BCAAs in all combinations suppressed the growth defect of *bat1*Δ cells, especially when valine was present (Figure 3B), in agreement with a proposal that valine synthesis in yeast is primarily a mitochondrial process regulated by Bat1 (Takpho *et al*, 2018). Supplementation with the corresponding alpha-keto acids did not suppress the growth defect of *bat1*Δ cells (Figure EV3).

**FIGURE 3.**
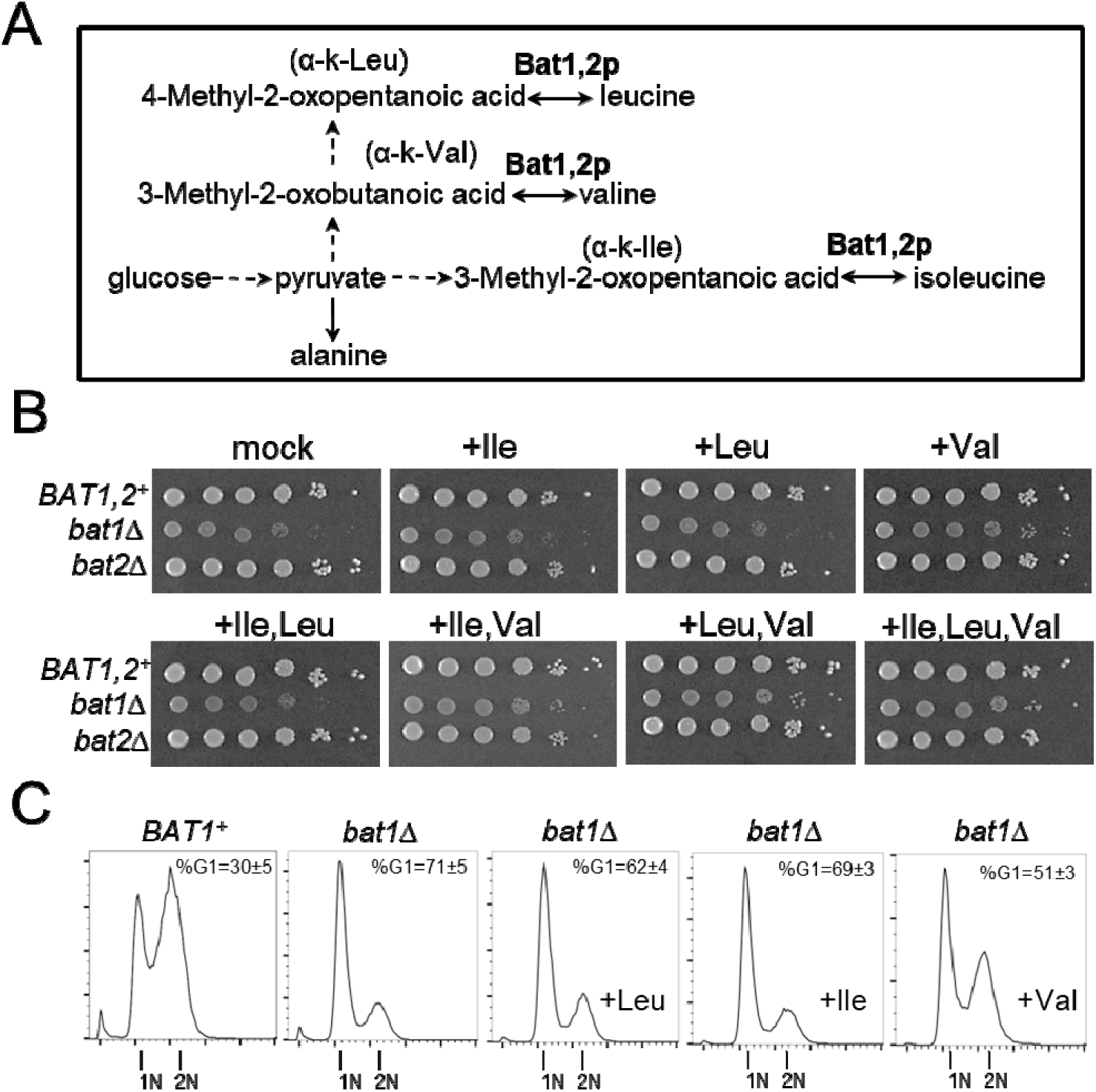
BCAA supplementation suppresses the growth defect of *bat1*Δ mutants. A, Diagram of the reactions leading to BCAAs from the corresponding alpha-keto (α-k) acids, catalyzed by Bat1,2p. A more detailed diagram leading to the of valine (M5) and leucine (M6) isotopomers is in Figure EV1. B, The indicated strains (all in the prototrophic CEN.PK background; see Materials and Methods) were spotted at 10-fold serial dilutions on solid Synthetic Minimal Medium (SMM) agar plates. Exogenous amino acids were added at 1mM final concentration, as indicated in each case. The plates were incubated at 30 °C and photographed after 3-days. C, DNA content profiles of *BAT1* and *bat1*Δ cells from asynchronous cultures, in SMM medium. Where indicated, exogenous amino acids were added at 1mM final concentration. On the x-axis of the histograms is fluorescence per cell, while on the y-axis is the cell number. Peaks corresponding to cells in G1 with unreplicated (1N) and cells in G2 and M phases with fully replicated (2N) DNA are indicated. The percentage of cells with G1 DNA content (%G1) from 3 independent measurements is shown in each case (mean and sd). The values used to generate the graphs are in File S1/Sheet3.

However, even when all three BCAAs were added exogenously, they did not fully restore the growth of *bat1*Δ cells (Figure 3B). There are several possibilities to explain these results: First, *de novo* BCAA synthesis by the Bat1 mitochondrial enzyme may be required for optimal growth. Second, Bat1 may have novel roles. It has been speculated that Bat1 may be enzymatically involved in synthesizing iron-sulfur clusters in mitochondria, which are then exported to the cytoplasm (Prohl *et al*, 2000), or even that Bat1 has an unknown non-enzymatic role (Kingsbury *et al*, 2015).

To link the poor growth of *bat1*Δ cells with specific metabolic alterations in these cells, we examined their steady-state metabolite levels using several approaches. First, we measured the intracellular levels of amino acids via the PTH-based detection method, as in Figure EV2. With this assay, the intracellular levels of isoleucine and, especially, valine were significantly lower in *bat1*Δ compared to *BAT1^+^* cells (Figure EV4). Interestingly, the levels of Gly and Ala were also lower in *bat1*Δ cells (Figure EV4). We next compared metabolite levels between *BAT1^+^*and *bat1*Δ cells using two different MS-based approaches. Primary metabolites were measured with GC-TOF MS, while biogenic amines with HILIC-QTOF MS/MS (see Materials and Methods). Using mass spectrometry, the levels of all three BCAAs, including leucine, were significantly lower (p<0.05) in cells lacking Bat1 (Figure EV5, FileS1/Sheet10). Overall, however, from the >300 compounds detected and assigned from the mass spectrograms, the levels of only 15 compounds changed significantly by >1.5-fold between *BAT1^+^* and *bat1*Δ cells (Figure EV5, FileS1/Sheet10). Some dipeptides were enriched in either *BAT1^+^* or *bat1*Δ cells (p<0.0001 with Holm-Bonferroni correction, based on the MetaboAnalyst platform (Chong *et al*, 2019)), while the ‘amino acids’ metabolite group was enriched in *BAT1^+^* vs. *bat1*Δ cells (p<0.02 with Holm-Bonferroni correction, based on the MetaboAnalyst platform). Alterations in TCA cycle metabolites have been reported for double *bat1,2*Δ cells (Kingsbury *et al*, 2015), but these changes were not evident in the single, prototrophic *bat1*Δ cells we evaluated here. In summary, the above results suggest that a drop in intracellular BCAAs, and not widespread global metabolite changes, is at least in part responsible for the slow growth of *bat1*Δ cells.

### Bat1-deficient cells are small and delayed in the G1 phase of the cell cycle

To see how the slow growth of *bat1*Δ cells manifests during cell division, we first looked at the DNA content of asynchronously dividing *BAT1^+^* and *bat1*Δ cells. Asynchronous *bat1*Δ cultures had at least twice as many cells in the G1 phase than *BAT1^+^* cells (Figure 3C). Exogenously added BCAAs (at 1mM final concentration) had varying effects. Isoleucine did not modify the DNA content profile, but leucine and valine reduced the fraction of cells with 1N DNA content (Figure 3C). The effect was much more substantial for valine (51% of cells with 1N DNA content, vs. 71% for mock-treated cells) but still far from the distribution seen in wild type cells (30% of cells with 1N DNA content; compare the left histogram to the right one in Figure 3C). These results mirror the results from the growth assays in Figure 3B, arguing that exogenous BCAAs, especially valine, suppress the phenotypes of *bat1*Δ cells, but they do not fully restore their growth and cell cycle progression.

We next examined cell cycle-associated phenotypes in more detail. In budding yeast, changes in the kinetics of cell cycle progression are often accompanied by cell size changes. As nutrient availability and quality are reduced, the birth and mean size of cells are reduced, the rate at which cells increase in size is lowered, and the duration of the G1 phase increases disproportionately compared to the duration of subsequent cell cycle phases (Johnston *et al*, 1977; Soma *et al*, 2014). Loss of Bat1 significantly reduced both the birth and mean cell size in asynchronous cultures in SMM medium (Figure EV6A). We then used centrifugal elutriation to obtain highly synchronous, early G1 daughter cell populations. Wild type and *bat2*Δ cells progressed in the cell cycle with similar kinetics (Figure EV6B, compare the left and right panels). They had a critical size (defined as the size at which half the cells are budded) of ∼37 fL. On the other hand, the slow growth of *bat1*Δ cells made elutriation very challenging, and only about 60% of the elutriated cells in the population eventually budded (Figure EV6B, middle panel). The small elutriated *bat1*Δ cells increased in size very slowly, having a specific rate of size increase (*k*) of 0.15h^-1^ vs. 0.23h^-1^ for wild type cells (Figure EV6C). From all these measurements, knowing the birth size (Vb), the size at which cells enter the S phase (critical size, Vc), and the rate at which they increase in size (*k*), we can obtain accurate estimates of the absolute length of the G1 phase (T_G1_), from the equation T_G1_=(LN(Vc/Vb))/*k*. While *BAT1^+^* daughter cells stay in G1 for ∼2.3 h, in *bat1*Δ daughters, the G1 phase lasts ∼4.5 h. We conclude that *bat1*Δ cells have a pronounced G1 delay, resembling nutrient-limited cells.

### Bat1 is functionally linked to TOR activation

It has been previously reported that TOR signaling is significantly reduced in loss-of-function *bat* mutants in yeast (Kingsbury *et al*, 2015). It is possible then that the growth defect of *bat1*Δ cells could at least in part reflect TOR inhibition. Since rapamycin inhibits the TOR complex 1 (TORC1), which drives cellular anabolic processes (Loewith & Hall, 2011), we queried the growth of *bat1*Δ cells in the presence of sub-lethal levels of rapamycin (Figure 4A). The growth of *BAT1^+^* and *bat1*Δ cells was equalized by rapamycin added at 30 ng/mL (Figure 4A). These results suggest that the inhibition of TORC1 by rapamycin lies downstream of Bat1, and it is epistatic to the growth defects of *bat1*Δ cells. To further test this notion, we looked at a molecular output of TORC1 signaling. Phosphorylation of ribosomal protein S6 (Rps6) reports on TORC1 activation, and it can be detected in yeast using an antibody against the human phosphorylated S6 protein (Wallace *et al*, 2022; González *et al*, 2015). While phosphorylated Rps6 was readily detected in wild-type cells in minimal medium (Figure 4B), it was barely so in *bat1*Δ cells (Figure 4B, top blot after prolonged exposure). We confirmed that the signal attributed to Rps6 phosphorylation is sensitive to TORC1 inhibition by rapamycin (Figure 4C, left) and phosphatase treatment (Figure 4C, right).

**FIGURE 4.**
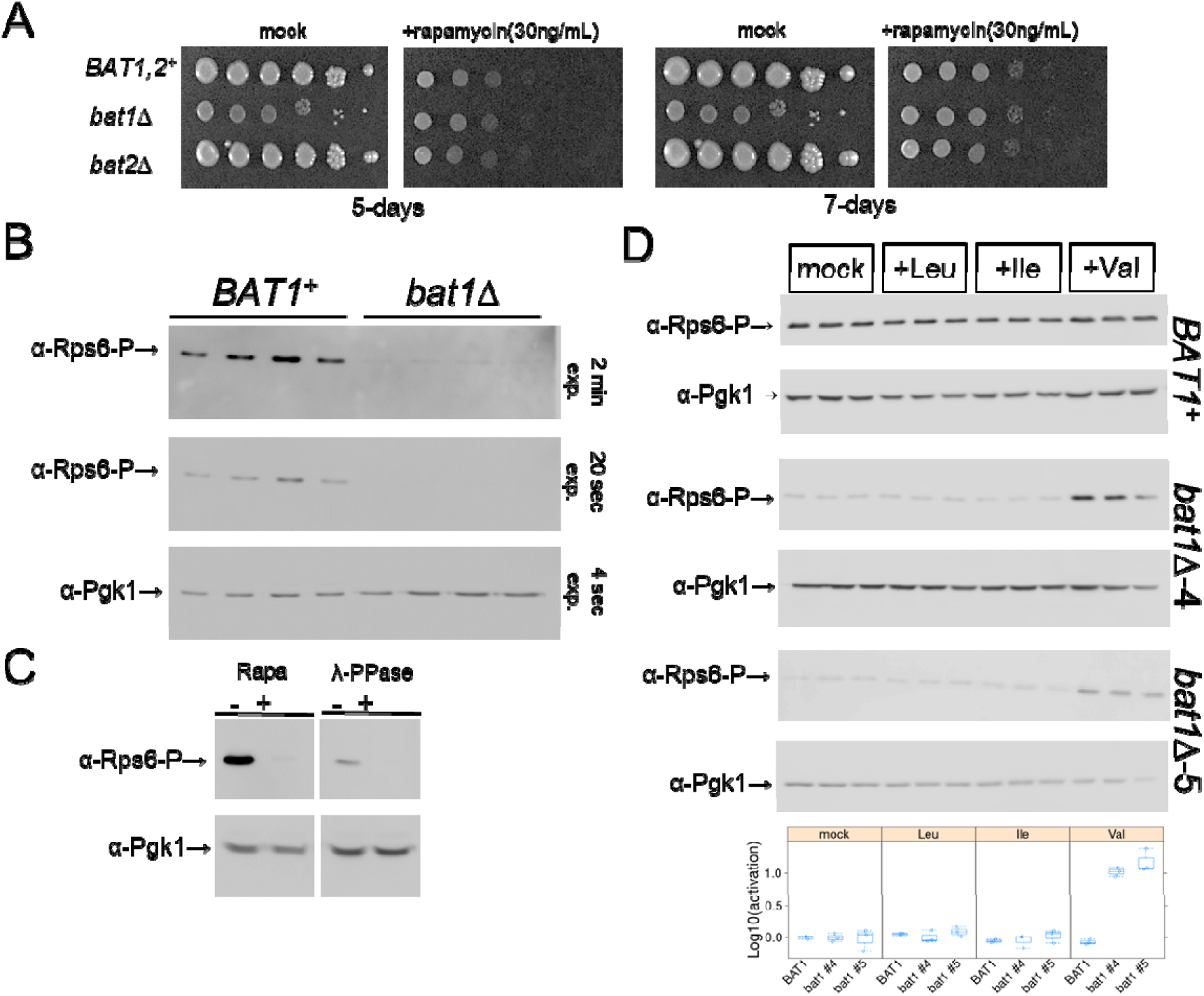
Bat1 is functionally linked to TORC1 activation in early G1. A, The indicated strains were spotted at 10-fold serial dilutions on solid Synthetic Minimal Medium (SMM) agar plates. Rapamycin was added at 30 ng/mL, as shown in each case. The plates were incubated at 30 °C and photographed after 5 and 7 days, as indicated. B, Immunoblots of total cell extracts from asynchronous *BAT1* and *bat1*Δ cells, from four independent experiments. The signal from Pgk1 (α-Pgk1) is shown on the blot at the bottom, and from phosphorylated Rps6 (α-Rps6-P) is on the blot above, indicated for two exposures (2 min, top; 20 sec, middle). C, On the left are immunoblots from wild-type (CEN.PK) cells treated (+), or not (-), with rapamycin (Rapa) at 200ng/mL for 1 h before cell extract preparation. On the right are immunoblots of wild-type (CEN.PK) cell extracts. The extracts were treated (+), or not (-), with λ-phosphatase for 1 h (see Materials and Methods). The levels of phosphorylated Rps6 (α-Rps6-P) and Pgk1 (α-Pgk1) are shown in each case. The raw immunoblots for B & C are in File S5. D, Exogenous addition of valine leads to sustained activation of TORC1 and phosphorylation of Rps6 in cells lacking Bat1. Wild type (*BAT1^+^*) and bat1Δ (two independent isolates, #4, and #5) strains were grown overnight in minimal (SMM) medium, diluted to 1E+06 cells/mL in fresh SMM media containing the indicated amino acid (at 1mM), and harvested when they reached 5E+06 cells/mL. The levels of phosphorylated Rps6 (α-Rps6-P) and Pgk1 (α-Pgk1) are shown in immunoblots from total cell extracts in each case. The relative levels of Rps6-P/Pgk1 are shown in each case at the bottom. The values used to generate the graphs are in File S1/Sheet4. The raw immunoblots for D are in File S6.

Since the exogenous addition of BCAAs suppressed the growth defect of *bat1*Δ cells (Figure 3B), albeit to varying degrees, we next measured the levels of phosphorylated Rps6 upon adding BCAAs. For this experiment, overnight cultures of *BAT1+* and *bat1*Δ cells were diluted to the same cell density (1E+06 cells/mL) in fresh minimal media containing the indicated amino acid (at 1mM) and harvested after they had completed at least two cell divisions when they reached 5E+06 cells/mL. In wild type cells, the magnitude of elevated levels of phosphorylated Rps6 by exogenous BCAAs was minimal (e.g., only ∼10% in the case of leucine (Figure 4D)). This agrees with other reports, which showed that adding BCAAs does not lead to sustained TORC1 activation in wild type cells (Kingsbury *et al*, 2015; Stracka *et al*, 2014). On the other hand, we found that *bat1*Δ cells exposed to valine had >10-fold higher phosphorylated Rps6 (Figure 4D). These results were evident in two independently derived *bat1*Δ strains (*bat1*Δ*-4* and *bat1*Δ*-5*; Figure 4D), mirroring the ability of exogenous valine to suppress the slow growth of *bat1*Δ cells (Figure 3B,C) and their reduced TORC1 activity (Figure 4B). We note that exogenous valine has also been reported previously to activate TORC1 in some settings (Mirisola *et al*, 2014). Overall, the most straightforward interpretation of our results is that cells lacking Bat1 may represent a highly sensitized background of low TORC1 activity, enabling the display of effects (e.g., activation of TORC1 by exogenous BCAAs) that are difficult to detect in prototrophic wild type cells with high TORC1 activity.

TORC1 activity already is at a maximal level in wild type cells cultured with a preferred nitrogen (ammonium) and carbon (glucose) source (Hughes Hallett *et al*, 2014), making it difficult to enhance it further. It is mostly in the context of loss-of-function perturbations (e.g., TORC1 inhibition by rapamycin or nutrient withdrawal (Barbet *et al*, 1996; Loewith & Hall, 2011; Heitman *et al*, 1991; De Virgilio & Loewith, 2006; Moreno-Torres *et al*, 2015; Pedruzzi *et al*, 2003; Kingsbury *et al*, 2015; Hughes Hallett *et al*, 2014), or upon Bat1 loss (Figures 3,4, EV6)) that proliferative effects become apparent.

### TORC1 activity during cell division in wild type cells

Since we detected an increase in leucine synthesis as cells progress in the cell cycle in wild-type cells (Figure 2), demonstrated a severe G1 delay (Figures 3 and EV6) and reduced TORC1 activity (Figure 4) in *bat1*Δ cells, we then examined TORC1 activity in the cell cycle in wild-type cells. It has been known for decades that inhibition of TORC1 by rapamycin arrests budding yeast (Heitman *et al*, 1991) and mammalian (Albers *et al*, 1993) cells in the G1 phase of the cell cycle. But just because inhibition of TORC1 by genetic or chemical means leads to G1 arrest is not enough to conclude that in dividing, unperturbed cells, there is dynamic control of TORC1 activity in the cell cycle. On the contrary, one could argue that since bulk cell growth and protein synthesis appear to have a constant rate throughout the cell cycle in constant nutrients in budding yeast (Elliott & McLaughlin, 1978), perhaps TORC1 activity is also constant in these conditions. We had previously shown that the abundance of the Rps6 paralogs, Rps6A or Rps6B, or any other ribosomal protein, does not change at all in the cell cycle (Blank *et al*, 2020). Here, we found that Rps6 phosphorylation rises steadily in synchronous cultures of wild-type prototrophic cells in minimal medium, increasing ∼3-4-fold by the time cells have exited the G1 phase when they are large (>35 fL) and budded (>50%) (Figure 5A, bottom blots; and 5B, for the quantification from multiple experiments). These results provide evidence of cell cycle-dependent changes in TOR signaling in budding yeast.

**FIGURE 5.**
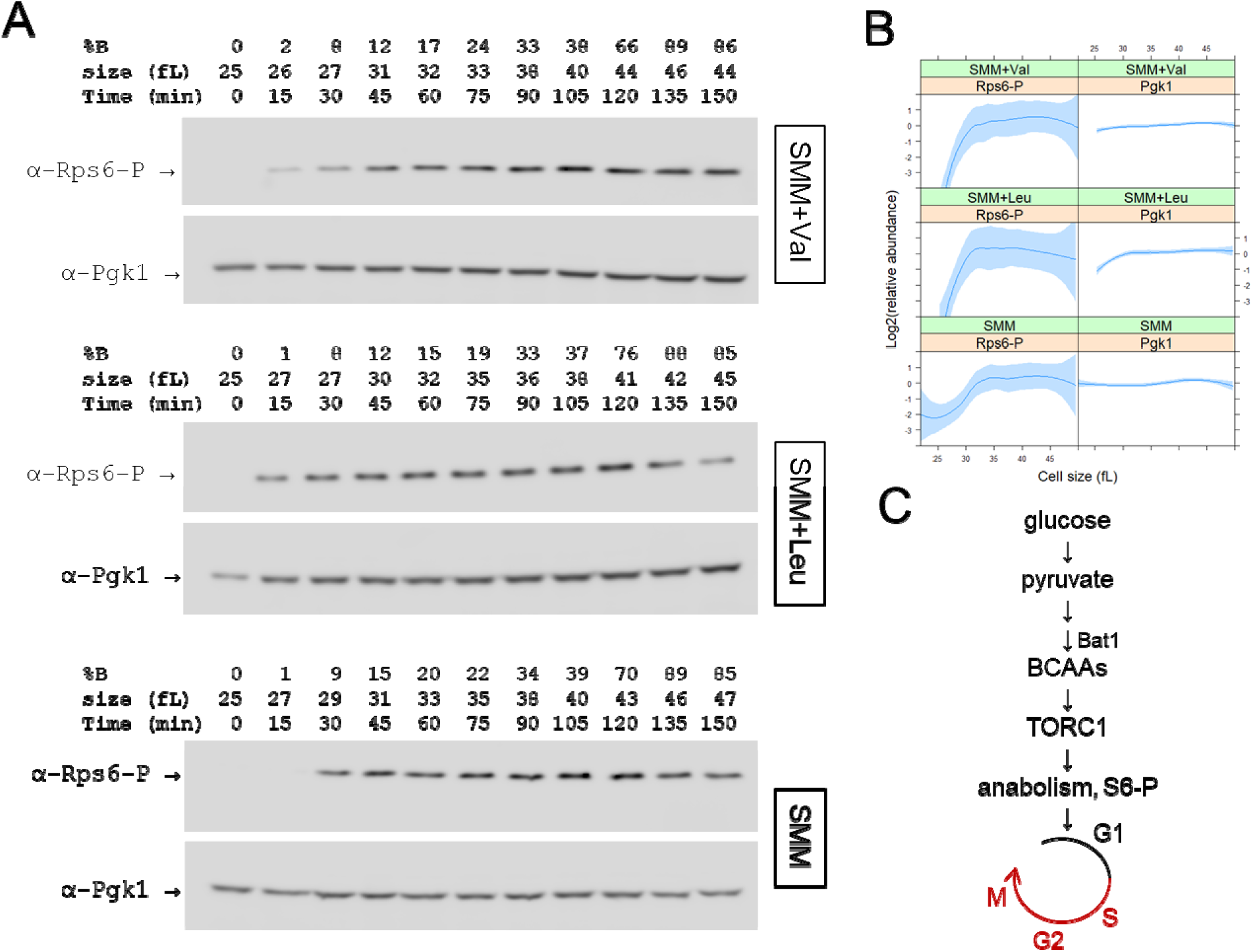
TORC1 activity increases as cells progress in the cell cycle. A, Immunoblots of total cell extracts from synchronous, elutriated wild-type (CEN.PK) cells in minimal (SMM) medium with exogenous Leu or Val added at 1mM immediately after elutriation. At the top, the percent of budded cells (%B), cell size (in fL), and time (in min) are indicated. The levels of phosphorylated Rps6 (α-Rps6-P) and Pgk1 (α-Pgk1) are shown in each case. B, Quantification of the levels of phosphorylated Rps6 and Pgk1 from independent experiments done as in A (see Files S7,8). The relative levels of each protein across the cell cycle is shown on the y-axis, as a function of cell size (x-axis). Loess curves and the std errors at a 0.95 level are shown. The values used to draw the plots are in File S1/Sheet5. C, Schematic of a possible model to explain our findings, linking BCAA synthesis to TORC1 activation early in the cell cycle.

Although exogenous BCAAs did not lead to a sustained increase in TORC1 activity in asynchronously proliferating wild type cells (Figure 4D), we next asked if they might change the pattern of the temporal activation of TORC1 in the cell cycle. We isolated newborn daughter cells in early G1 phase, and resuspended them in media supplemented with 1mM leucine (Figure 5A, middle blots) or valine (Figure 5A, top blots). A small acceleration of the rise in the levels of phosphorylated Rps6 was evident in both the leucine-and valine supplemented cells (Figure 5A,B). Nonetheless, there were no noticeable downstream consequences in the kinetics of cell cycle progression, in either the rate the cells increased in size or their critical size (Figure 5A; see values above the corresponding blots), consistent with the notion that TORC1 activity already is at a maximal level in these conditions (Hughes Hallett *et al*, 2014).

## Conclusions

The experiments we described provide the first picture of changing BCAA synthesis in synchronous, proliferating yeast cells. Together, they point to a straightforward sequence of events, from glycolytic flux and generation of pyruvate, to BCAA synthesis by the Bat1 aminotransferase, and then TORC1 activation late in G1 and into the S phase (Figure 5C). In developing eye imaginal discs in flies, TORC1 activity scored by phosphorylation of Rps6 has also been reported to be the highest in S phase cells (Kim *et al*, 2017; Romero-Pozuelo *et al*, 2017). Our isotope tracing results argue that despite the apparent constancy of overall cell growth, specific pathways of central metabolism, such as leucine biosynthesis, may show cell cycle-dependent changes (Figure 2). This view of dynamic, cell cycle-dependent control of metabolic pathways is consistent with recent findings in budding yeast using single-cell microscopy, showing that ribosome biogenesis (Guerra *et al*, 2022) and protein synthesis (Takhaveev *et al*, 2023), known to be regulated by TORC1 and protein kinase A pathways, change in the cell cycle, peaking in G1 and then later also in G2/M. Lipid biosynthesis also strongly depends on cell cycle progression in yeast, peaking in the G2 and M phases (Blank *et al*, 2020; Takhaveev *et al*, 2023). As the assays develop further, both with isotope tracing and single-cell approaches, more biosynthetic pathways will likely be found to display dynamic temporal control during cell division. Such advances may revise the long-held view of cellular biosynthesis as a static, ‘housekeeping’ process with few prominent landmarks during cell proliferation.

It is unknown how the activity of Bat1 may be regulated. The steady-state levels of the *BAT1* transcript (Spellman *et al*, 1998; Blank *et al*, 2020, 2017), and Bat1 protein (Blank *et al*, 2020), do not change in the cell cycle. On the other hand, Bat1 is heavily modified post-translationally at multiple residues by phosphorylation (Lanz *et al*, 2021; Albuquerque *et al*, 2008; Zhou *et al*, 2021), acetylation (Henriksen *et al*, 2012), and succinylation (Frankovsky *et al*, 2021; Weinert *et al*, 2013), which could affect its enzymatic activity. Lastly, we note that the mammalian ortholog BCAT1 is activated and necessary for chronic myeloid leukemia (CML) development in humans and mice, exercising its oncogenic function through BCAA production in CML cells (Hattori *et al*, 2017). Interestingly, the therapeutic potential of rapamycin in CML was the objective of a recently completed clinical trial (NCT00780104), but the results are not available yet. Future work, perhaps in CML cell lines, incorporating synchronous cultures and isotope tracing, may test if our yeast results extend to mammals, including humans.

## MATERIALS AND METHODS

A Reagent Table is in the Supplementary Files. Where known, the Research Resource Identifiers (RRIDs) are shown in File S9.

### Strains and media

All the strains used in this study are shown in the Reagent Table. Unless indicated otherwise, the cells were cultivated in the standard synthetic minimal media (SMM), containing 0.17% ^w^/_v_ yeast nitrogen base without amino acids and ammonium sulfate, 0.5% ^w^/_v_ ammonium sulfate, 2% ^w^/_v_ glucose at 30°C (Kaiser *et al*, 1994). To generate the *bat1*Δ haploid strain, we used the single-step gene replacement method of Longtine (Longtine *et al*, 1998). Briefly, a PCR product was generated using plasmid pFA6a-kanMX6 as a template (Longtine *et al*, 1998), with primers BAT1-F1 and BAT1-R1, and used to transform strain CEN.PK. Positive colonies resistant to G418 (Geneticin) were identified and genotyped by PCR with primers VIII:517387-FWD and VIII:518942-REV, to confirm that *BAT1* was absent and replaced by the kanMX marker. Similarly, using primers BAT2-F1 and BAT2-R1, we generated the *bat2*Δ strain and genotyped it with primers X:705443-FWD and X:706939-REV. During strain construction, the cells were grown in agar plates with the standard rich undefined medium YPD (1% ^w^/_v_ yeast extract, 2% ^w^/_v_ peptone, 2% ^w^/_v_ dextrose) containing G418 at 100 μg/mL.

### Centrifugal elutriation, cell size and DNA content measurements

General methods have been described previously (Hoose *et al*, 2012; Soma *et al*, 2014). Briefly, for each elutriation we used ∼250 mL of culture at 1-2E+07 cells/mL, in SMM medium. The cells were loaded on the elutriator at a pump speed of 35 mL/min and centrifuge speed of 3200 rpm. The cells were washed with 250 mL at 2800 rpm, and another 250 mL at 2400 rpm. They were then elutriated with 250 mL at a pump speed of 38 mL/min and centrifuge speed of 2400 rpm. For the isotope tracing experiments, the elutriated early G1 cells were split into eight samples and allowed to proceed in the cell cycle in a medium with ^12^C-glucose for varying times, as indicated in Figure 1A. Then, to label newly synthesized metabolites, we switched the medium to one with uniformly labeled ^13^C-glucose, and the cells were cultured for a 20 min ‘pulse’ period. The culture was then switched back to a ^12^C-glucose medium for a 20 min ‘chase’ period, and the cells were harvested for metabolite extraction (He *et al*, 2014). The experiment was repeated four more times from independent elutriation experiments, generating separate pulse-chase time courses. Cell size and budding measurements from these experiments are in File S1/Sheet1.

To accurately quantify the percentage of cells with 1N DNA content (%G1; see Figure 3C), we first measured the total area of the histogram. Then, we measured the area of the left half of the 1N peak, multiplied it by two, and divided it by the total area value. This fraction was multiplied by 100 to obtain the %G1 values shown in each case (Figure 3C and File S1/Sheet3).

### Metabolite extraction for isotope tracing

The lyophilized cell extracts were first quenched by adding to the sample microcentrifuge tubes 0.4 ml methanol (kept at –20 °C) and 400 μl double-distilled water (ddH_2_O) with ^13^C-glutaric acid (1μg/ml, 4 °C) used as internal standard. Then, 400 μl chloroform (at –20 °C) was added, and the samples were mixed on a rotating shaker (at 1400 rpm) for 20 min at 4 °C. The samples were centrifuged for 5 min at 16,000 *g* at 4 °C, and 300 μl of the supernatant were transferred to GC glass vials (Klaus Trott Chromatographiezubehör, Germany) and dried under vacuum at 4 °C overnight (Labconco, United States). The vials were then capped with magnetic caps (Klaus Ziemer GmbH, Germany) and stored at –80 °C until measurements.

Before derivatizing the supernatant samples, 50 μl of a ^13^C glutaric acid solution (10 μg/ml) in ddH_2_O was added as an internal standard to each sample and dried under vacuum as described above.

### Derivatization and GC-MS measurements

The sample derivatization was performed by adding 30 μl 2% methoxyamine hydrochloride (MeOX) in pyridine (Roth, Germany) for 90 min at 55 °C, followed by the addition of 30 µl N-tert-butyldimethylsilyl-N-methyltrifluoroacetamide (MTBSTFA) and incubated for 60 min at 55 °C. The tert-butyldimethylsilyl (TBDMS) derivatized samples were analyzed with a gas chromatograph connected to a mass spectrometer (GC-MS) (Agilent 7890A and 5975C inert XL Mass Selective Detector). For each measurement, 1 µl of the sample was injected in splitless mode into an SSL injector at 270 °C. The GC was equipped with a 30 m DB-35MS + 5 m Duraguard capillary column (0.25 mm inner diameter, 0.25 µm film thickness). Helium was used as carrier gas at a flow rate of 1.0 mL/min, and the GC-oven was run at the following temperatures and times per sample: 6 min 80 °C; 6 min 80 to 300 °C; 10 min 300 °C; 2.5 min 300 to 325 °C; 4 min 325 °C. Each GC-MS run lasted 60 minutes. The transmission temperature from GC to MS was 280 °C, and the MS and the quadrupole temperatures were 230 °C and 150 °C, respectively. The ionization in the mass detector was performed at 70 eV. The detector was operated in SCAN-mode for full scan spectra from 70 to 800 m/z with 2 scans/s. We used a C10-C40 alkane standard mixture (Sigma-Aldrich) to calibrate retention indices.

### Isotope tracing data analysis

Data analysis was performed with the Metabolite Detector (Hiller *et al*, 2009), RStudio, and LibreOffice software packages. After calibration with the alkane mixture, a normalization by the internal standard ^13^C-Ribitol was performed to eliminate variations in sample volumes. Batch quantifications were performed with Metabolite Detector. Non-targeted analysis was performed with an in-house library and the following settings: Peak threshold 5; minimum peak height 5; 10 bins per scan; deconvolution with 5; no baseline adjustment; compound reproducibility 1.00; maximum peak discrimination index 100.00; min # ions 20.

All the data from these experiments are in File S2. The MID values from the cell extracts, and the associated p-values from ANOVA analysis to detect significant differences, are in File S2/Sheet1, and File S2/Sheet2, respectively. Similarly presented are the data for the MID values from culture supernatants (File S2/Sheet3 and File S2/Sheet4), and the SIM data from cell extracts (File S2/Sheet5 and File S2/Sheet6) and culture supernatants (File S2/Sheet7 and File S2/Sheet8). To normalize and display the data in the cell cycle as shown in Figure 2, we processed the values as described previously (Spellman *et al*, 1998), yielding their “relative abundance” in the cell cycle. Briefly, the value of a species at any one time point in a cell cycle series was divided by the average of all values of that species at all time points in that series. These normalized values were then Log2-transformed and used to generate the graphs in Figure 2.

### Statistical analysis, sample-size and replicates

For sample-size estimation, no explicit power analysis was used. All the replicates in every experiment shown were biological ones, from independent cultures. A minimum of three biological replicates was analyzed in each case, as indicated in each corresponding figure’s legend. The robust bootstrap ANOVA was used to compare different populations via the t1waybt function, and the posthoc tests via the mcppb20 function, of the WRS2 R language package (Wilcox, 2011; Mair & Wilcox, 2020). The non-parametric Kruskal-Wallis and posthoc Nemenyi tests were done with the posthoc.kruskal.nemenyi.test function of the PMCMR R language package. No data or outliers were excluded from any analysis.

### Immunoblot analysis

For protein surveillance, protein extracts were made as described (Wallace *et al*, 2022), and resolved on 12% Tris-Glycine SDS-PAGE gels, unless indicated otherwise. Loading was measured with an anti-Pgk1p primary antibody (at 1:5,000; ThermoFisher; Cat#: 459250), followed by a secondary antibody (at 1:5,000; ThermoFisher; Cat#: 31431). Ribosomal protein S6 phosphorylation was detected by a specific rabbit monoclonal antibody against Ser235/236 of the human protein (at 1:5,000; Cell Signaling, Cat#: 4858), followed by a secondary antibody (at 1:5,000; ThermoFisher; Cat#: 31466). Imaging and quantification was done as described previously (Blank *et al*, 2020, 2017; Maitra *et al*, 2020, 2022). Signal intensities in the cell cycle (see Figure 5A) are shown as Log2-transformed relative abundance ratios, as described above (Spellman *et al*, 1998).

### PTH-based amino acid analysis

The HPLC-based measurements of PTH-derivatized amino acid levels was done at the Protein Chemistry Laboratory of the Department of Biochemistry and Biophysics, as described previously (Maitra *et al*, 2020; He *et al*, 2014).

### Steady-state metabolite profiling

The untargeted, primary metabolite and biogenic amine analyses were done at the NIH-funded West Coast Metabolomics Center at the University of California at Davis, according to their established mass spectrometry protocols, from 1-2E+07 cells, as described previously (Blank *et al*, 2020; Maitra *et al*, 2020, 2022). Primary metabolites were measured with GC-TOF MS, while biogenic amines with HILIC-QTOF MS/MS. The raw data for the primary metabolite measurements are in File S3. The raw data for the primary amine measurements are in File S4. To identify significant differences in the comparisons among the different strains, we used the robust bootstrap ANOVA, as described above. The input values we used were first scaled-normalized for input intensities per sample. Detected species that could not be assigned to any compound were excluded from the analysis.

### Data availability

Strains are available upon request. The authors affirm that all data necessary for confirming the conclusions of the article are present within the article, figures, and tables. The Expanded View, Supplementary Material and Source data includes the following:

Figure EV1: Diagram showing the carbon additions and eliminations from glucose (M6) to the synthesis of valine (M5) and leucine (M6) isotopomers.

Figure EV2: Steady-state amino acid levels in the cell cycle from PTH-based analysis.

Figure EV3: Alpha-keto acid supplementation does not suppress the growth defect of *bat1*Δ mutants.

Figure EV4: Amino acid composition of *BAT1^+^* and *bat1*Δ strains from PTH-based analysis.

Figure EV5: Volcano plot of steady state abundances of primary metabolites and biogenic amines.

Figure EV6: Cells lacking Bat1 are smaller, increase in size slower, and are delayed in the G1 phase of the cell cycle.

File S1: Source data for the graphs in all Figures.

File S2: Metabolite measurements from the GC-MS measurements during the cell cycle, from the isotope tracing experiments.

File S3: Steady-state primary metabolite measurements in *BAT1^+^*vs. *bat1*Δ cells.

File S4: Steady-state biogenic amine measurements in *BAT1^+^* vs.*bat1*Δ cells.

File S5: Source immunoblots for Figures 4B,C.

File S6: Source immunoblots for Figure 4D. File S7: Source immunoblots for Figures 5A,B.

File S8: Additional source immunoblots for Figure 5B. File S9: Reagent Table

## Supporting information

Source data for the graphs in all Figures

Metabolite measurements from the GC-MS measurements during the cell cycle, from the isotope tracing experiments

Source immunoblots for Figures 4B,C

Source immunoblots for Figure 4D

Source immunoblots for Figures 5A,B

Additional source immunoblots for Figure 5B

Reagent Table

Point-by-point author response

Marked-up file with text changes

## ACKNOWLEDGEMENTS

We thank Maria Cardenas-Corona (NIH) for reagents. This work was supported by the National Institutes of Health (NIH, grant R01 GM123139) and by the German Research Foundation (DFG, Project-ID 34509606—TRR 51).

## AUTHOR CONTRIBUTIONS

M.P. and K.H. conceived and designed the study. C.R. and K.S-H. performed and analyzed the isotope tracing experiments. S.E.H. performed the cell cycle analyses shown in Figure 3C. H.B. and M.P. performed all other experiments. All authors evaluated and discussed the data. M.P., H.B., and C.R. wrote the first draft of the manuscript, which was then edited by all authors.

## EXPANDED VIEW FIGURES

**FIGURE EV1.**
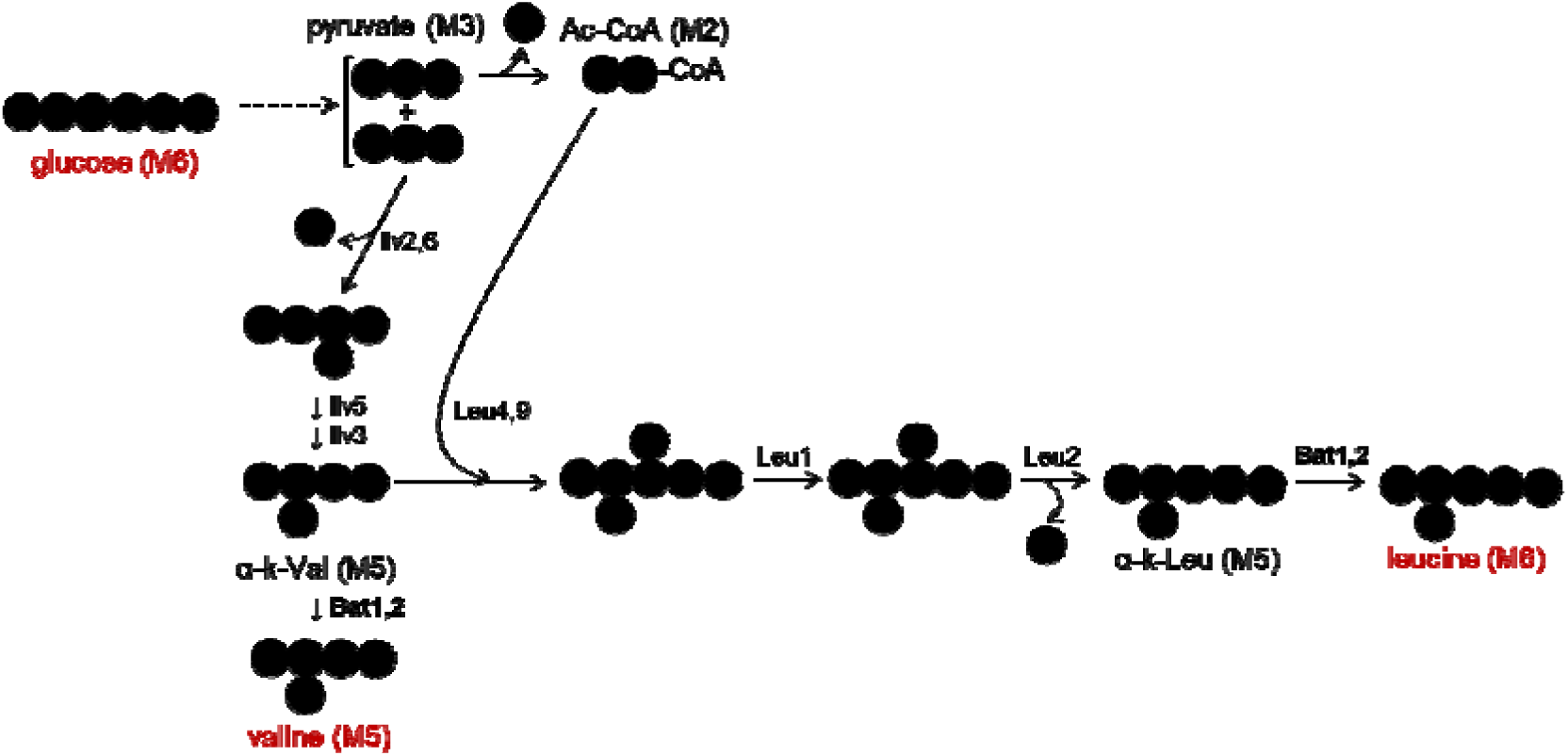
Diagram showing the carbon additions and eliminations from glucose (M6) to the synthesis of valine (M5) and leucine (M6) isotopomers.

**FIGURE EV2.**
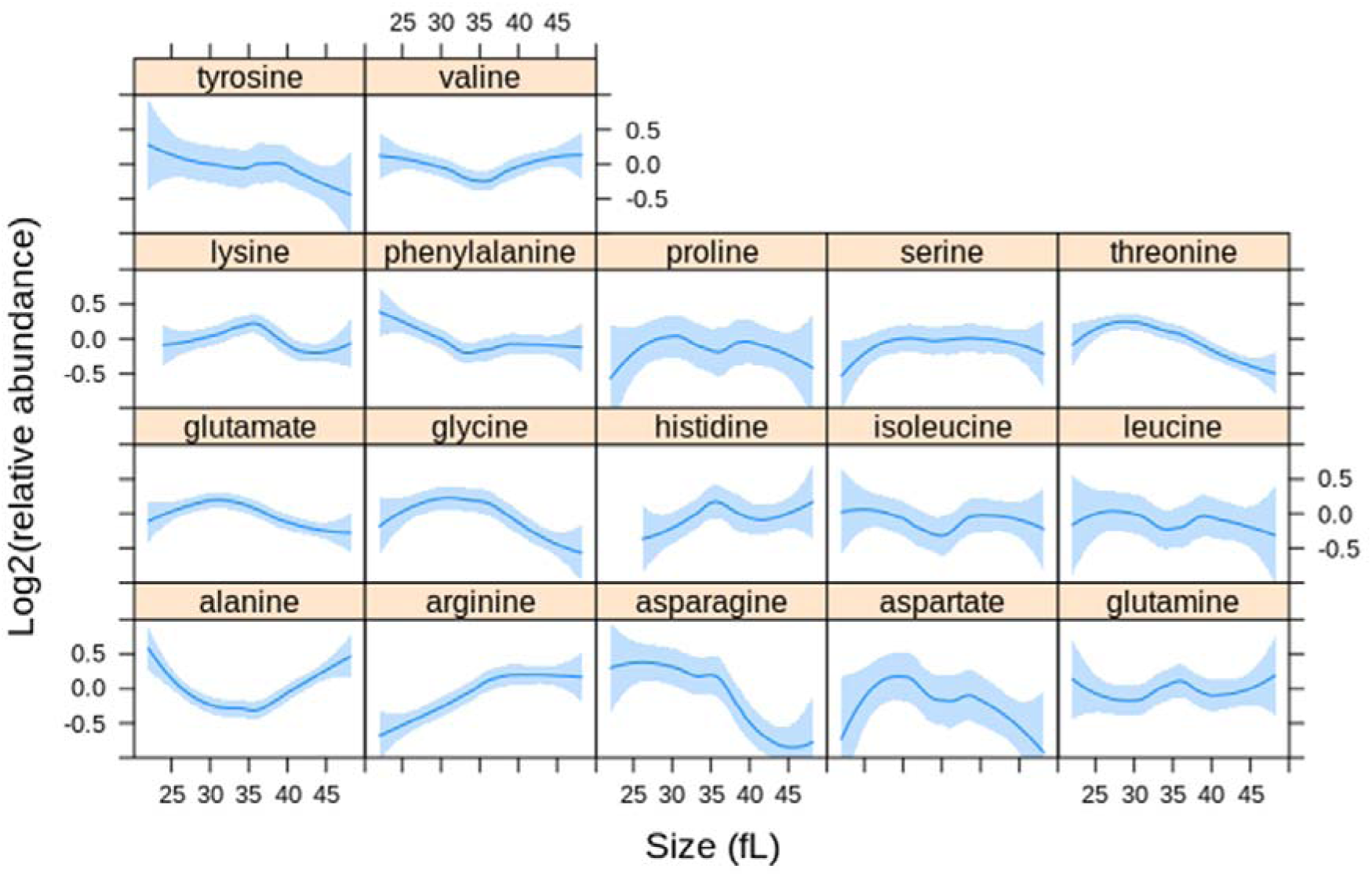
Steady-state amino acid levels in the cell cycle. Intracellular amino acid levels were measured as described in Materials and Methods from synchronous, elutriated cultures in SMM media. The fraction of each amino acid is on the y-axis. The values used to draw the plots are in File S1/Sheet6.

**FIGURE EV3.**
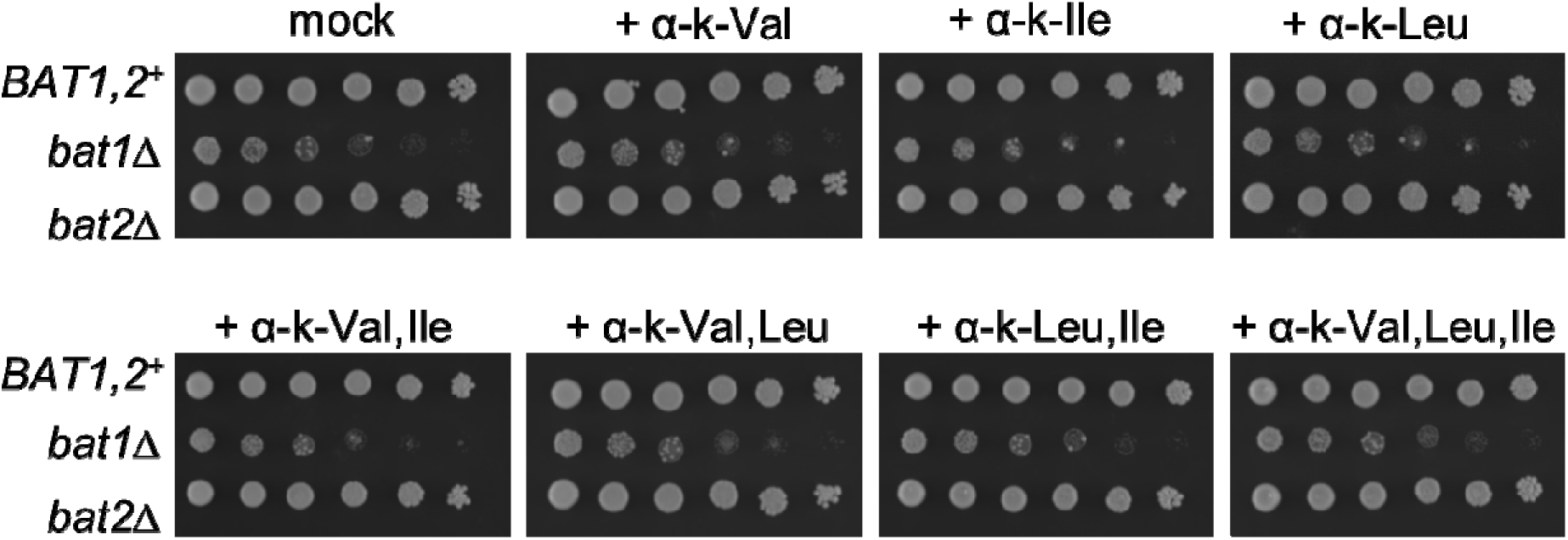
Alpha-keto acid supplementation does not suppress the growth defect of *bat1*Δ mutants. The experiment was done as in Figure 3. Exogenous alpha-keto acids were added at 1mM final concentration, as indicated in each case.

**FIGURE EV4.**
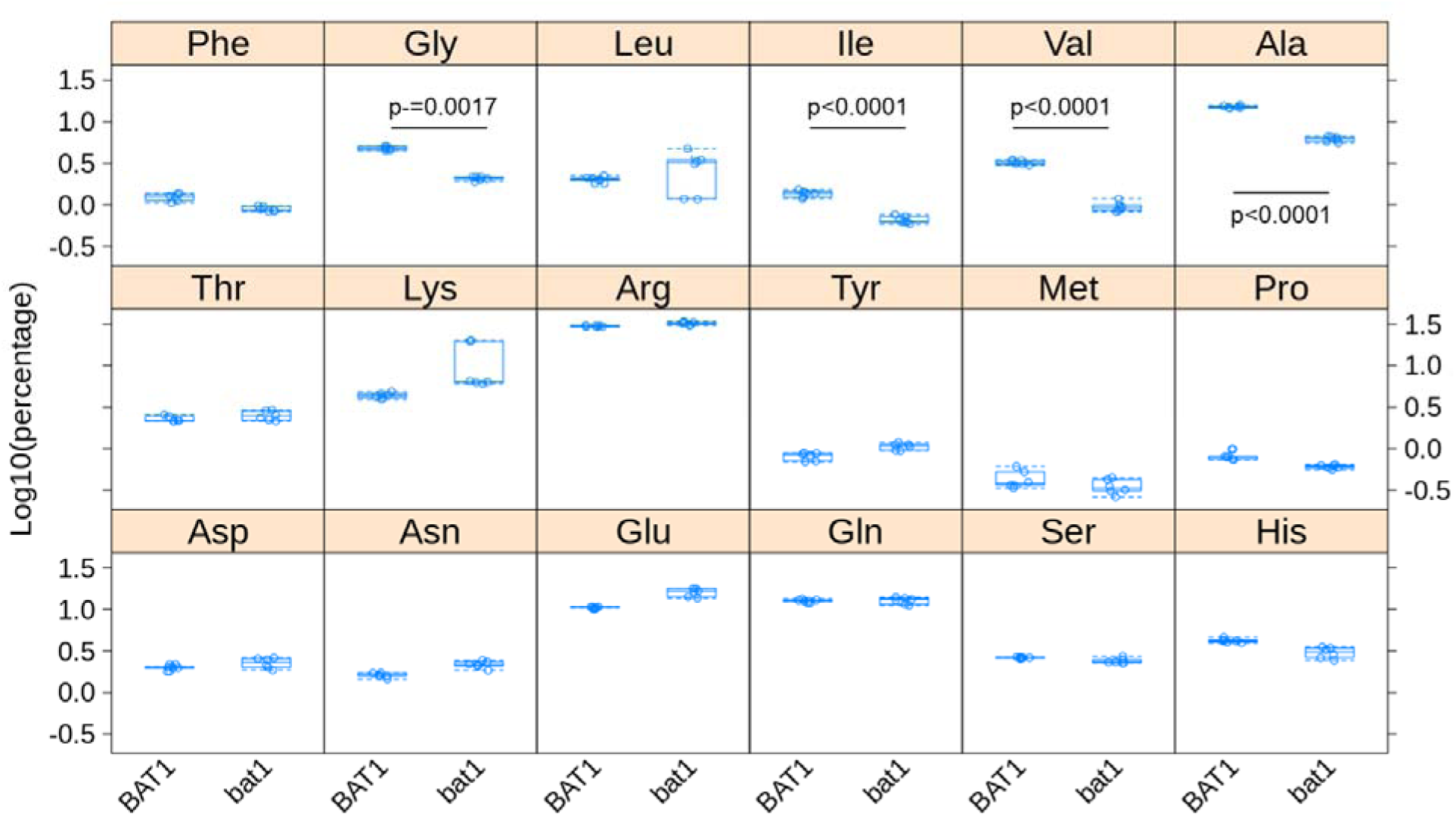
Amino acid levels in *BAT1^+^* and *bat1*Δ cells. Intracellular amino acid levels were measured as described in Materials and Methods and Figure EV2, from exponentially growing cultures in SMM media. The fraction of each amino acid is on the y-axis. The four amino acids (Gly, Ile, Val, Ala) with significantly altered fractional abundance (>2-fold, p<0.05; from six independent samples in each case) are indicated. The values used to draw the plots are in File S1/Sheet7.

**FIGURE EV5.**
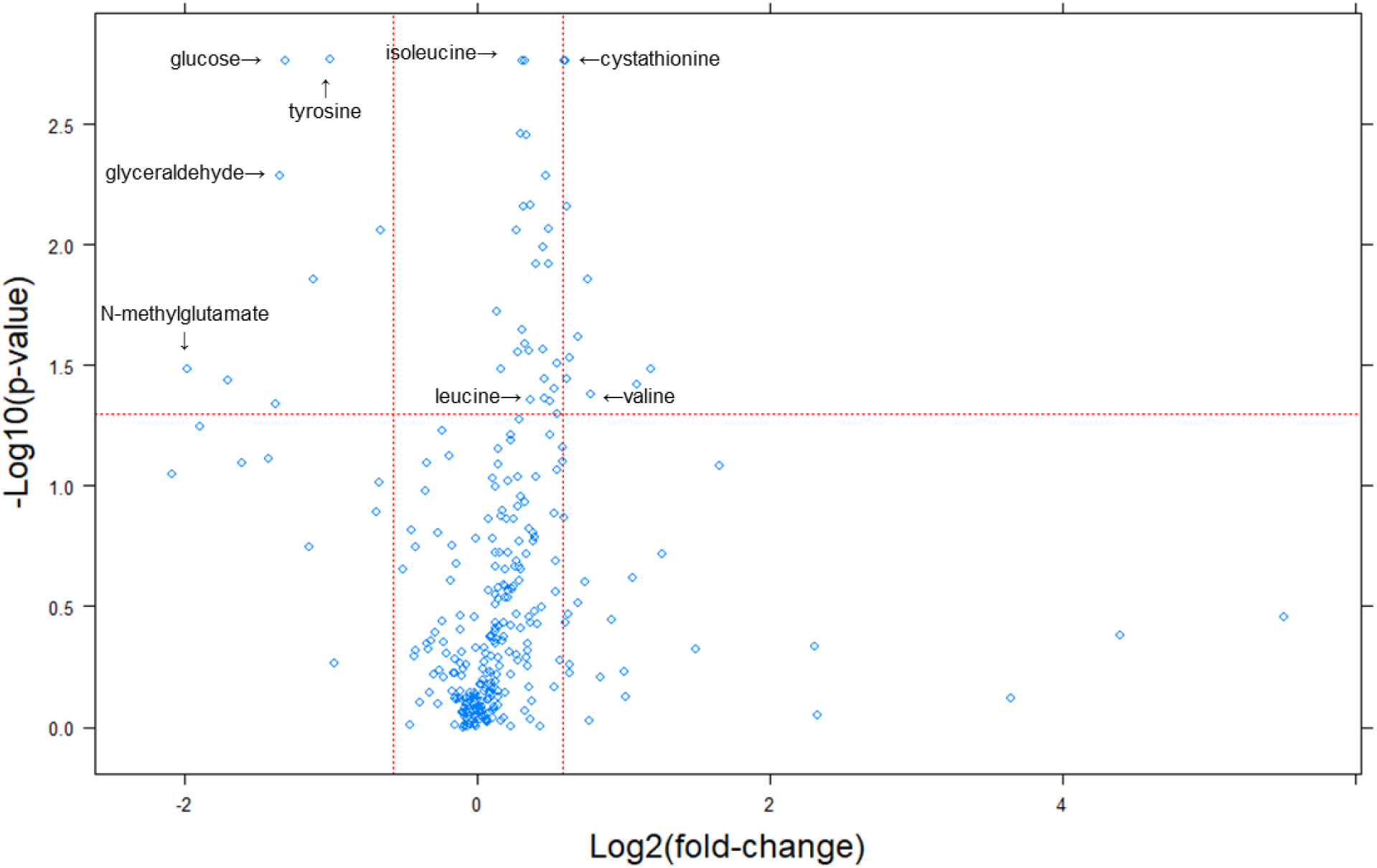
Changes in primary and biogenic amine metabolites in cells lacking Bat1. Metabolites whose levels changed were identified from the magnitude of the difference (x-axis; Log2-fold change in *BAT1^+^*: *bat1*Δ cells) and statistical significance (y-axis), indicated by the red lines. The analytical and statistical approaches are described in Materials and Methods. The values used to generate the graphs are in File S1/Sheet8. The labels indicate selected metabolites.

**FIGURE EV6.**
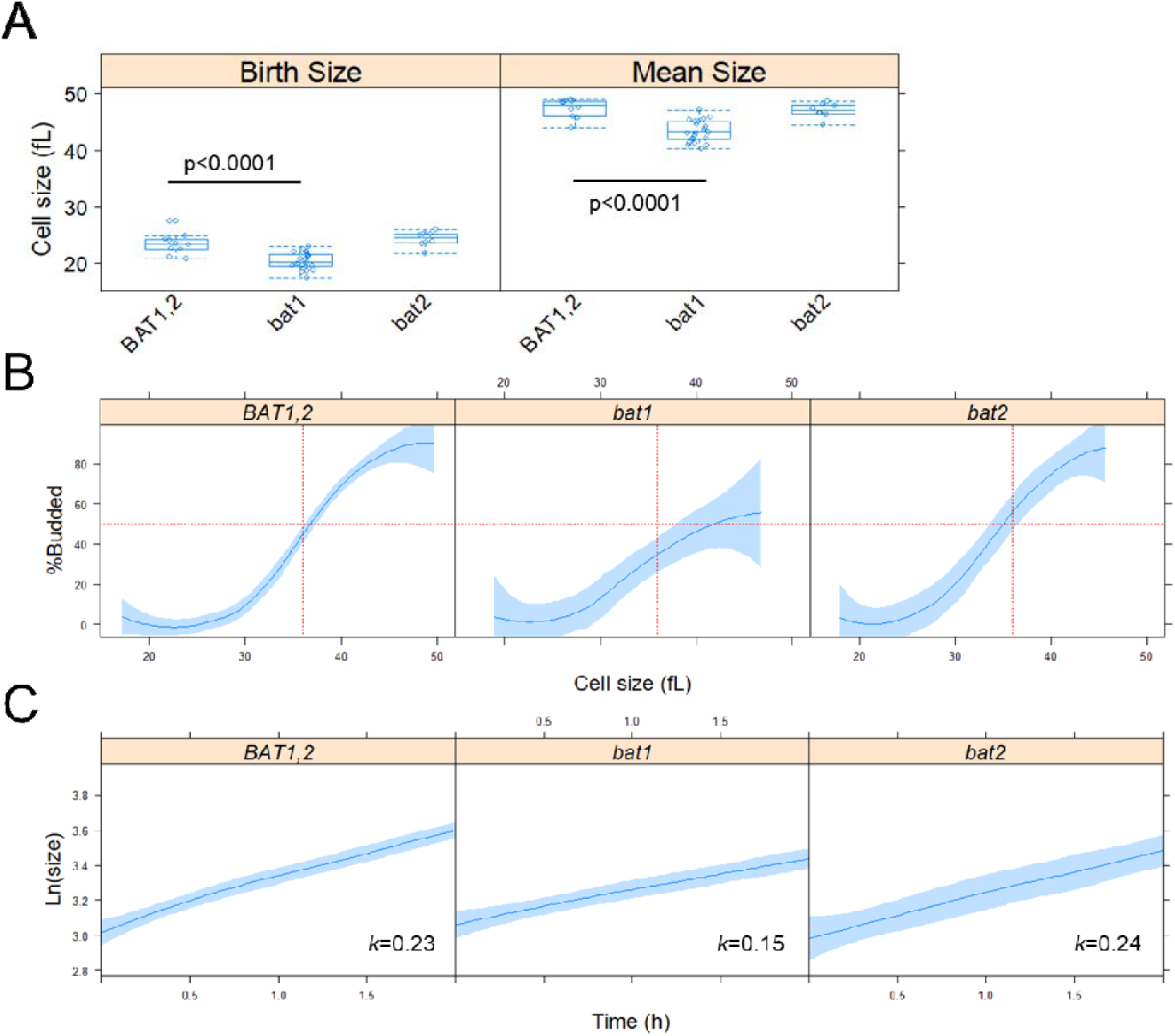
Cells lacking Bat1 are smaller, increase in size slower, and are delayed in the G1 phase of the cell cycle. A. Boxplots showing the size of the cells (in fL). The measurements were taken from asynchronous cultures in synthetic minimal media (SMM) without amino acid supplementation. The values used to draw the plots are in File S1/Sheet9. B, Plots of the percentage of budded cells (y-axis) as a function of size (x-axis). The measurements were from daughter cells of the indicated strains, obtained by centrifugal elutriation, progressing in the cell cycle in SMM medium. Loess curves and the std errors at a 0.95 level are shown. C, From the same experiments as in B, the specific rate of the increase in size (*k*) was calculated, from the slope of the regression lines plotting the Ln-transformed cell size values (y-axis) against time (x-axis). The values used to draw the plots in B and C are in File S1/Sheet10.

