## Supplementary figures and images for "Branched chain amino acid synthesis is coupled to TOR activation early in the cell cycle in yeast"

### Source immunoblots for Figure 4D

#### File S6. Source data immunoblots related to Figure 4D

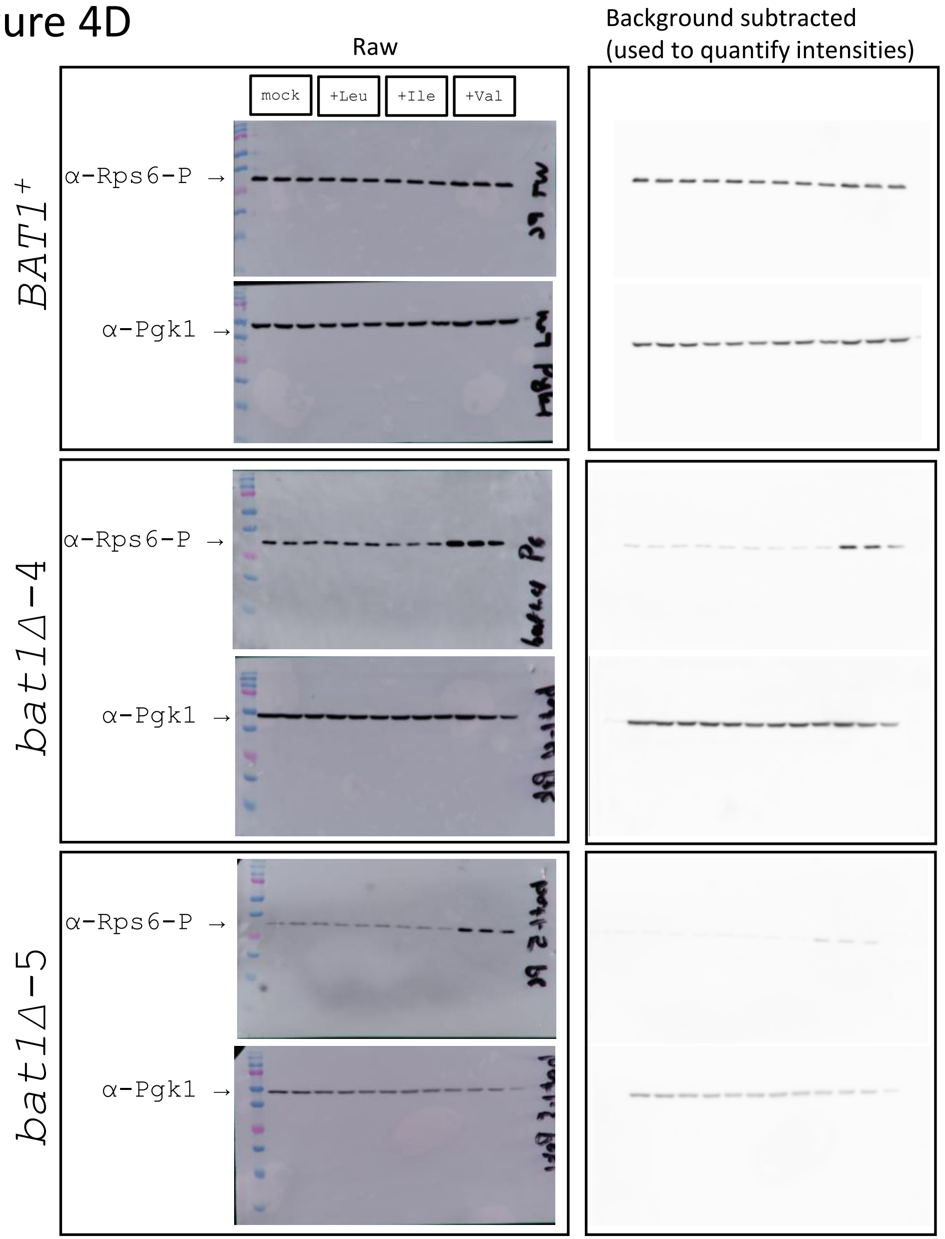

### Source immunoblots for Figures 4B,C

#### File S5. Source data immunoblots related to Figure 4B & 5C

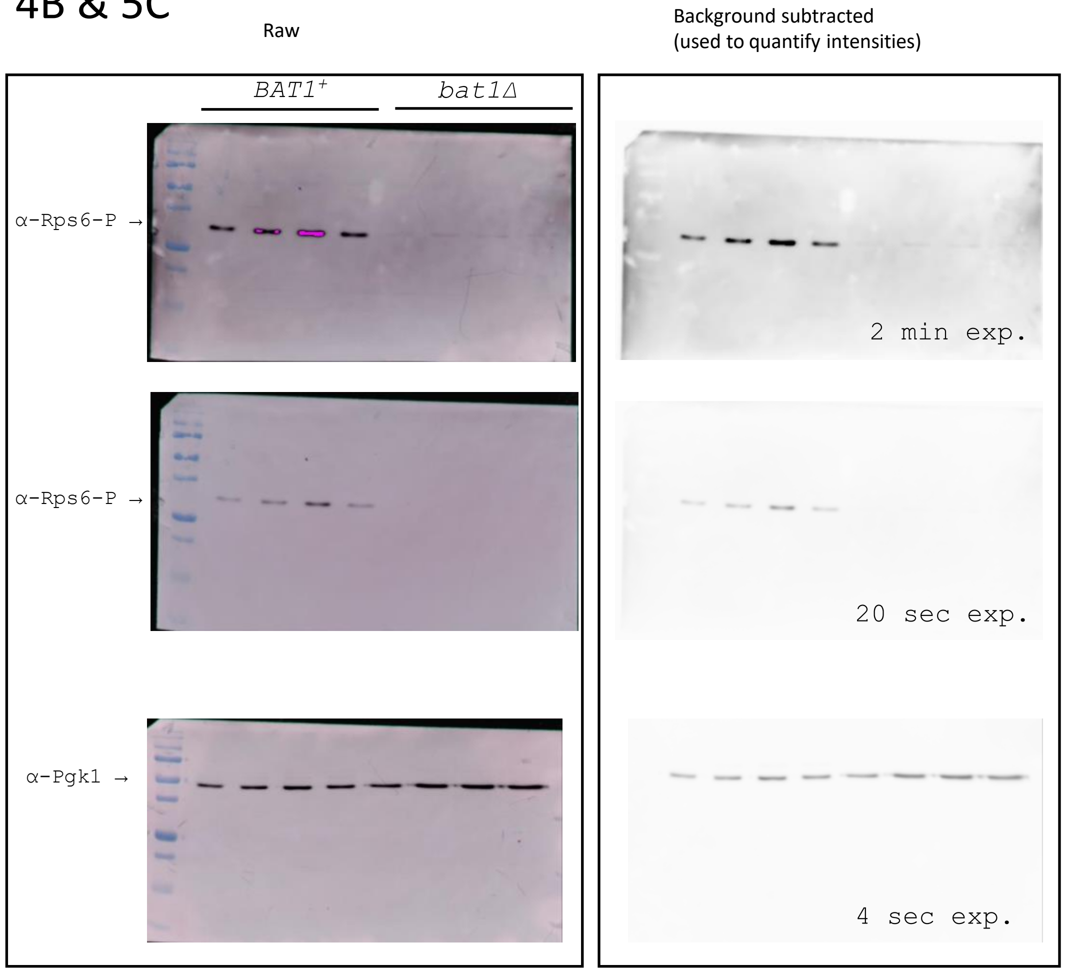


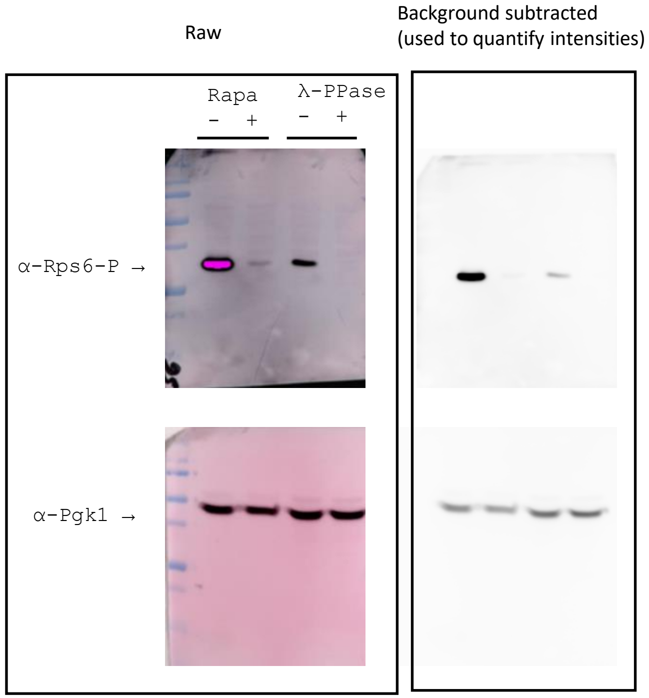

### Source immunoblots for Figures 5A,B

#### File S7. Source data immunoblots related to Figure 5A & 5B

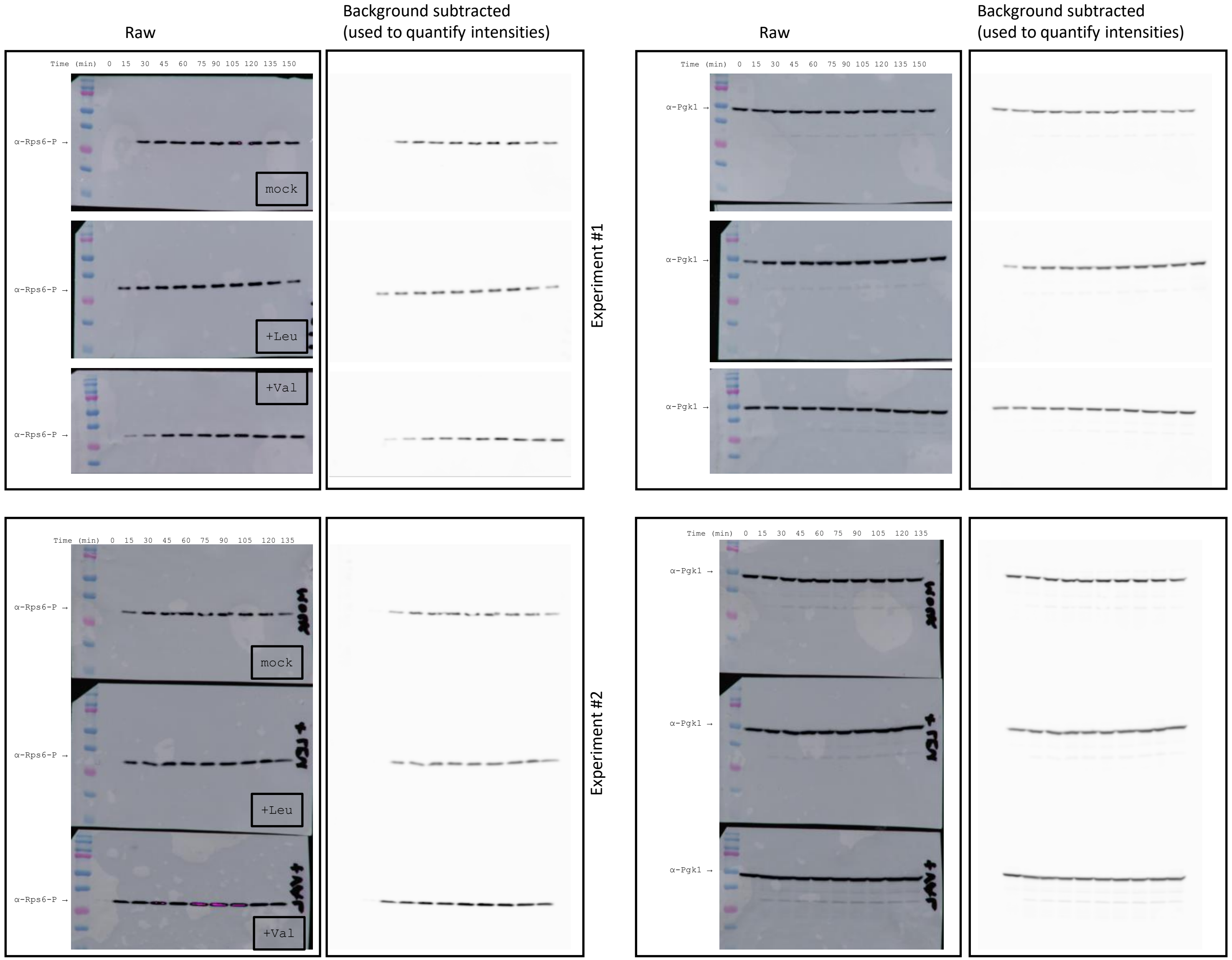
