## Additional source immunoblots for Figure 5B for "Branched chain amino acid synthesis is coupled to TOR activation early in the cell cycle in yeast"

### File S8. Source data immunoblots for additional elutriated wild type (CEN.PK) cells related to Figure 5B

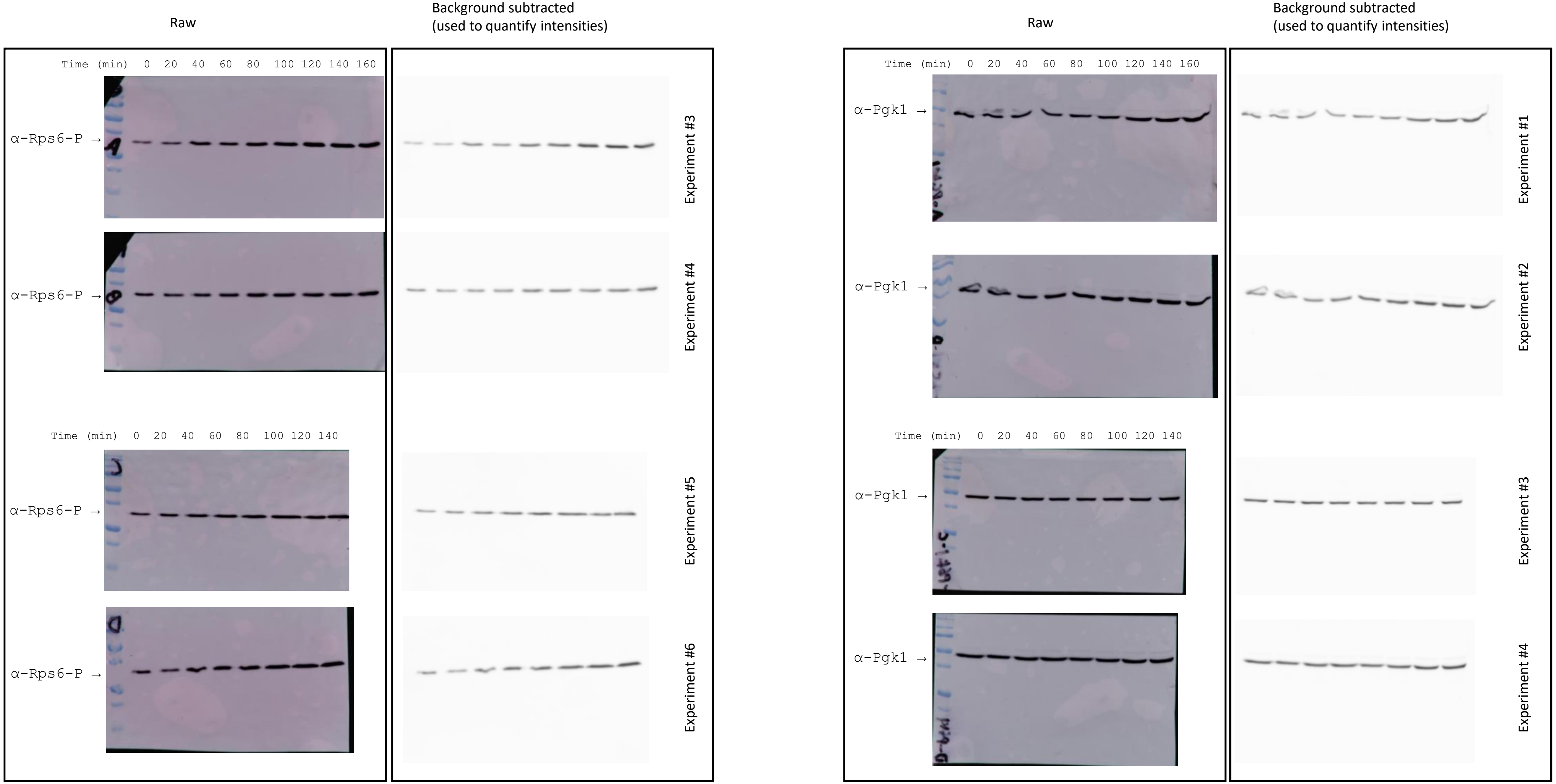
