## Supplementary material for "Branched chain amino acid synthesis is coupled to TOR activation early in the cell cycle in yeast": Point-by-point author response

Full Revision
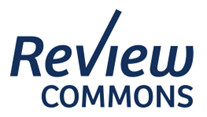

**Manuscript number: RC-2023-01825**

**Corresponding author(s): Michael Polymenis and Karsten Hiller**

1. **General Statements**

We thank all reviewers for their comments and suggestions. The revised manuscript included new experiments they suggested and extensive text edits. Our point-by-point response is shown in bold.

**2.	Point-by-point description of the revisions**

—----------------------------------------------------------------------------------------------------------------

Reviewer #1 (Evidence, reproducibility and clarity (Required)):

Summary

In this manuscript, Blank et al. propose a link between cell-cycle dependent changes in metabolic flux and corresponding changes in TORC1 activity in yeast cells. Based on their findings, the authors propose that Bat1-dependent leucine synthesis from glucose increases as cells progress through G1 and that this activates TORC1 to drive cell cycle progression. Although the existence of cell-cycle dependent synthesis of leucine is a novel and exciting finding, several aspects of the proposed model are not sufficiently supported by experimental evidence, in particular the fact that the increase in Leu synthesis is causing the increase in TORC1 activity in late G1.

Major comments:

1. To show that the increase in Leu biosynthesis in S-phase is activating TOR, one would ideally want to blunt this increase in biosynthesis and assay TORC1 activity. Admittedly, this is difficult. So, instead, the authors study bat1- cells which have strongly impaired synthesis of BCAA including Leucine. The relevance of these bat1- cells to the proposed cell-cycle dependent model, however, is questionable for two reasons: 1) Although the authors state that "exogenous supplementation of BCAAs in all combinations suppressed the growth defect of bat1- cells, especially when valine was present", the spot assays in Figure 3 show visible rescues only when valine is present either alone or in combination, while supplementation of leucine or isoleucine does not seem to have any effect. Hence it appears that the bat1- phenotype is mainly due to limiting valine levels, not leucine levels. 2) The relevance of these results for understanding TORC1 regulation are questionable, since valine does not typically activate TORC1. Does addition of Leu to bat1- cells increase TORC1 activity ?

**RESPONSE: The reviewer’s comments were very valuable. We performed the suggested experiments (adding not only Leu but also Ile and Val) to bat1 cells and measuring phosphorylation of Rps6 (see new Figure 4D) and the DNA content of those cells (see new Figure 3C). We found that Leu weakly promotes cell cycle progression, compared to the addition of Val, which also leads to pronounced activation of TORC1 (>10-fold activation; see Figure 4D). We discuss these findings in the revised text.**

**We also note, as published by others and now discussed in the text, that in WT cells, exogenous addition of Leu (or any other BCAA) does not lead to sustained activation of TORC1 (see new Figure 4D). This is not surprising. As reported by the Hall lab (see PMID: 25063813, which we now cite), the Gtr-dependent activation of TORC1 by BCAAs mentioned by the reviewer is very transient. Hence, our new data, showing sustained TORC1 activation and cell cycle effects upon Val addition in bat1 cells, is exciting. They argue that bat1 cells serve as a highly sensitized background of low TORC1 activity, enabling the display of effects that are difficult to measure in WT cells.**

2. TORC1 activity is known to depend on steady-state leucine concentrations in the cell rather than on leucine flux. Although the authors observe that the synthesis rate of leucine increases during G1 progression, this does not necessarily translate into increased leucine concentrations in the cell. To support the claim that the increase in TORC1 activity during G1 progression depends on leucine, the authors would need to show that, not only leucine synthesis, but also overall leucine levels in the cell increase during G1 progression.

**RESPONSE: We did this experiment and now report the data (see new Figure EV2), using the Edman degradation-based assay. We found that changes in the steady-state levels of BCAAs had a similar pattern, and those changes were most significant for valine (rising 30-40% from late G1 to G2/M). Nonetheless, we note also that the kinetics of amino acid synthesis measured by our isotope tracing experiment need not match the steady-state levels of amino acids. Steady-state levels are affected by a multitude of parameters, only one of which is the rate of synthesis, as we now discuss in detail in the manuscript.**

3. To test whether the increase in Leu biosynthesis in S-phase activates TORC1, a few different approaches could be tested: 1) Since leucine activates TORC1 through the Gtr proteins, the authors could test whether rendering TORC1 resistant to low leucine through expression of constitutively active Gtrs abolishes the cell-cycle dependence in TORC1 activity. 2) Leu could be added to the medium of wildtype cells in G1 to the amount necessary to cause an increase in intracellular Leu levels similar to those seen in S-phase to test whether this increases TORC1 activity.

**RESPONSE: We did the suggested experiments, which are now shown in the new Figure 5. Leucine and valine accelerated the rise in TORC1 activity in G1. However, there were no noticeable downstream consequences in the kinetics of cell cycle progression. As we discuss in the text:**

**“A small acceleration of the rise in the levels of phosphorylated Rps6 was evident in both the leucine- and valine supplemented cells (Figure 5A,B). Nonetheless, there were no noticeable downstream consequences in the kinetics of cell cycle progression, in either the rate the cells increased in size or their critical size (Figure 5A; see values above the corresponding blots), consistent with the notion that TORC1 activity already is at a maximal level in these conditions…”**

4. In Fig 2B one sees that Leu biosynthesis peaks at 150min and then drops again. The p-RpS6 blot in Fig. 5D, however, only goes up to 140 min and shows that TORC1 activity increases up to 140 min, but it doesn't show timepoints beyond 150 min when Leu biosynthesis drops again, and hence one would expect TORC1 activity to drop. If TORC1 activity were to drop from 150min onwards, this would strengthen the correlation between Leu biosynthesis and TORC1 activity.

**RESPONSE: The reason for the drop in Figure 2 is trivial and does not affect the interpretation. As seen in Figure 1 (the experiment from which the data in Figure 2 are shown), by 180 min, the cells were entering a new cell cycle, evidenced by a reduction in cell size (Figure 1B) and in the fraction of budded cells (Figure 2B). At that point, there is a mix of mothers and daughters with very poor synchrony, making it impossible to conclude much about the drop in Leu synthesis (i.e., does it arise from the lack of new synthesis in mothers, daughters, or both?). In the experiment in Figure 5, the reviewer mentions (now those figures have moved to File S8 because we added more experiments in the figure) the experiment terminated when peak budding was reached, which was 140 min, within one cell cycle. Lastly, it is important to stress that every elutriation experiment is different. While the times are close, comparing various experiments on a time basis alone is inaccurate. Instead, the metric used in the field to compare different experiments is usually cell size, which we use in all other Figures except Figure 1 because, in that case, the experiment was a time-based, pulse-chase one.**

Minor concerns:

1. In Figure EV4, the authors should highlight some of the metabolites that are significantly changed, in particular the BCAA. The figure is not very informative as currently presented.

**RESPONSE: We have now labeled the BCAAs, and a few more metabolites as suggested (note the Figure is now EV5).**

2. Fig 2 - are "expressed ratios" the best term for metabolite levels? Unlike genes, where such heat maps are often used, the metabolites are not 'expressed'. How about 'relative metabolite level' instead?

**RESPONSE: Good point. The axis now reads “relative abundance”.**

3. Page 8: "We also measured the MID values from the media of the same cultures used to prepare the cell extracts." Where are these data? We don't see them in File S2?

**RESPONSE: The data are in File S2 (there are many ‘sheets’ in the file). In sheets 3,4 are the MID values and the analysis from metabolites in the media.**

4. Fig 4B - the x-axis labeling is missing for the bat1- cells

**RESPONSE: Corrected. Note that new DNA content measurements are now shown in Figure 3C.**

5. Although the authors state repeatedly that they show "for the first time in any system" that TORC1 activity is dynamic in the cell cycle, similar observations have already been made before, for instance showing high mTORC1 activity in the G1/S transition in the Drosophila wing disc or low mTORC1 activity during mitosis in mammalian cells (see PMIDs 28829944, 28829945, and 31733992). The text should be amended accordingly.

**RESPONSE: Thank you. Corrected.**

6. There are two entries for valine in File S1/Sheet8. Why?

**RESPONSE: The reason is that they were detected in both analytical pipelines (primary metabolites and biogenic amines; primary metabolites were measured with GC-TOF MS, while biogenic amines with HILIC-QTOF MS/MS), which were combined in the Table. We did not describe it adequately in the previous version. We do now, in the Methods. We also note that the raw data from each method are shown in the corresponding supplemental files. We combined them in the Table used in the Figure for display purposes. We also note that the amino acids were also measured by another method (PTH-based HPLC). Hopefully, the new edits in the Methods clarify these points.**

Reviewer #1 (Significance (Required)):

Significance

Despite the well-known effects of pharmacological or genetic manipulations of TORC1/mTORC1 on cell cycle progression, whether and how mTORC1 activity itself is physiologically coupled to cell cycle progression is still an insufficiently studied aspect. Hence this study provides an interesting link between cell-cycle dependent regulation of amino acid biosynthesis and TORC1 regulation. Importantly, the results of this study rely on centrifugal elutriation to obtain cell cycle synchronization, thus ruling out potential metabolic artifacts due to pharmacological methods. The observed changes in metabolic flux are therefore likely genuine and represent the major strength of the study. The major limitation is the lack of strong evidence supporting the notion that the increase in Leu biosynthesis at late G1 or S-phase is causing the increase in TORC1 activity.

The major advance is conceptual - that amino acid biosynthesis rates are cell-cycle dependent.

These results will be of interest to a broad audience of people studying the cell cycle, cell growth, TORC1 activity, cell metabolism and cancer.

Reviewer #2 (Evidence, reproducibility and clarity (Required)):

This paper provides evidence that branched chain amino acid (BCAA) in the G1 phase of the cell cycle, fueled by pyruvate generated by glucose catabolism activates cell growth and allows cells to reach the critical size required for entry into S phase by activation of TORC1 signaling. Previous work had indicated that Leucine supplementation of a bat1 bat2 mutant, lacking both enzymes that catalyze BCAA from the alpha-keto acid precursors and starved on minimal medium, led to TORC1 activation. This work is significant in suggesting that BCAA synthesis from glucose is responsible for a cyclic activation of TORC1 necessary for a normal rate of cell growth in the G1 phase of the cell cycle.

The study employs metabolic flux analysis of metabolites derived from glucose following a pulse-chase with different isotopes of glucose in synchronized early G1 cells (obtained by elutriation) throughout one cell cycle. They claim that the only compelling changes in metabolites observed as the cell cycle proceeds was a decline in pyruvate containing only one heavy 13C carbon atom and a corresponding increase in Leu (M6) with 6 heavy carbon atoms, which is interpreted to indicate Leu synthesis from pyruvate that begins in early G1 and peaks at mitosis. They show that a bat1 mutant exhibits a slow-growth phenotype that can be mitigated only by valine (although they infer similar effects for Leu and Ile that I find unconvincing) and they observed reductions in all three BCAAs in different experiments that measure steady amino acid levels in different ways (although the results are compelling only for Val). They go on to show evidence that the bat1 mutation reduces birth and mean cell size and leads to an increased proportion of G1 cells in asynchronous cultures, and they claim that bat1 cells take much longer than WT to achieve the same size found when a synchronized WT culture reaches 50% budding (although they don't show the data for this last point.) Interestingly, they find that deleting BAT1 suppresses sensitivity to the TORC1 inhibitor rapamycin (Rap), consistent with the idea that the bat1 mutation impairs TORC1 activity in the same manner as Rap and that BCAA are required to activate TORC1 in WT cells to the level that can be impaired by Rap, as summarized in the model in Fig. 5F. Consistent with this, they present evidence that the bat1 mutation reduces TORC1 signaling as judged by diminished Rps6 phosphorylation (although it was not shown that this effect could be reversed by Val addition). They also show that TORC1 signaling/Rps6-P increases as the cell cycle progresses using elutriated early G1 cells, suggesting that TORC1 activity is periodic in the cell cycle (although they don't establish this periodicity through a second cell cycle).

General critique:

The conclusion that BCAA synthesis from glucose is responsible for a cyclic activation of TORC1 necessary for a normal rate of cell growth in the G1 phase of the cell cycle is potentially of considerable significance. There are however a number of puzzling aspects of the data that seem to weaken this conclusion. As described in greater detail below, it is difficult to explain why only Leu is synthesized from glucose during the cell cycle, and why only Val shows a marked reduction in the bat1 mutant that appears to be responsible for the slow-growth phenotype. In addition, there are important controls lacking of showing that a Val supplement can suppress the G1 delay and reduction in TORC1 signaling in the bat1 mutant. In addition, the evidence that TORC1 activity is periodic in the cell cycle is lacking and it needs to be shown that Rps6-P levels are periodic through at least a second cell cycle.

Major comments:

-Why don't they observe synthesis of Ile and particularly Val in the metabolic flux experiment of Fig. 1, especially considering that only Val appears to be critically required for normal cell growth in the bat1 mutant based on the results in Fig. 3B?

**RESPONSE: We now show the actual plots and the errors of all the measurements in Figure 2 (instead of a heatmap we had shown before). Valine (M5) levels show a very similar trend to leucine (M6). The variance in the measurements was higher, though, and statistically, the valine changes were less significant. Hence, it was more appropriate to highlight the leucine changes. Lastly, the new DNA content data (Figure 3C) show an effect upon the addition of leucine, albeit less significant than that of valine addition.**

-The data in Fig. 3B do not show a convincing increase in growth of the bat1 mutant with addition of Leu and Ile; and the stimulation by Val alone seems identical to that seen with Val in combination with Leu and Ile. Thus, it appears that the slow-growth of the bat1 mutant results only from reduced Val levels, not all 3 BCAAs, which is at odds with their interpretation of the data.

**RESPONSE. As mentioned above, the effect of valine is more pronounced than leucine's, but leucine does have consequences, best shown in the DNA content analysis (new Figure 3C). We also note that valine alone is insufficient to suppress the growth and cell cycle defects of bat1 cells. The latest data we have added (see Figures 3 and 5) are consistent with the interpretation that at least some de novo synthesis of BCAAs in the cell may be needed, explaining why exogenous BCAAs, including valine, are unable to correct the defects of bat1 cells fully.**

-they claim to see reductions in all three BCAAs in the bat1 mutant; however, no significant reduction was found for Leu in Fig. EV3, and only Val was altered by the 1.5-fold cut-off imposed on the MS metabolomics data in Fig. EV4 (which could be appreciated only by an in-depth examination of the supplementary data in File S1-the Val, Leu, and Ile dots should be labeled in Fig. EV4). In addition, the reductions in Ala and Gly showin in Fig. EV3 were not found in the MS analysis of Fig. EV4. It needs to be acknowledged that the metabolomics data show a marked reduction in the bat1 mutant only for Valine with little or no change in Leucine levels. This result is difficult to explain with the simple models shown in Fig. 3A and 5F, which requires additional comment. The authors should acknowledge the much greater effect of the bat1 mutation on Val levels versus Leu and Ile, revealed both by measuring the levels of BCAAs in the mutant and comparing the BCAAs for rescuing the slow-growth of the mutant, and explain how this can be reconciled with the results in Fig. 2 where only Leu and not Val or Ile synthesis was detected.

**RESPONSE. The perceived discrepancy in the steady-state measurements could easily arise from the different analytical methods used in each case. The differences are less substantial than the reviewer implies. For steady-state measurements in BAT1 vs. bat1 cells, we used the PTH-based method (which only detects amino acids) and two different MS-based pipelines (which detect various metabolites). From the MS-based analyses, the drop for all BCAAs was statistically significant. Although the magnitude of the drop was greater for valine (about 60% for valine vs. ~30% for isoleucine and leucine). Why is this a problem?**

**As for the valine changes in the isotope tracing experiments, as we mentioned above, the trend for valine (M5) was similar to that of leucine (M6) (now, hopefully, that data is shown better in Figure 2). Furthermore, as we commented above (see response to Reviewer 1) and now stated in the text, our isotope tracing experiments measure only the rate of synthesis, which need not match the steady-state abundances. The latter are affected by a multitude of variables, including the turnover of proteins and amino acids, not to mention their partition into distinct intracellular pools.**

**Lastly, please note that we have now added PTH-based measurements of amino acid levels in the cell cycle of wild type cells (new Figure EV2). As mentioned in our response to Reviewer 1, we found that changes in the steady-state levels of BCAAs had a similar pattern, and those changes were most significant for valine (rising 30-40% from late G1 to G2/M).**

-They need to add the data indicating that the bat1 mutant requires longer than WT cells to reach the ~35 fL volume at which 50% of WT cells are budded.

**RESPONSE: We added all that data (new Figure EV6) and discussed it better in the text. Note that our elutriation analyses allow accurate estimates of the G1 duration, which is at least 2x longer in bat1 vs. BAT1 cells.**

-It seems important to show that Val supplementation can suppress the overabundance of G1 cells in bat1 mutant cells shown in Fig. 4C; and can restore sensitivity to Rap and Rps6-P accumulation in bat1 mutant cells (in Fig.s 5A & B).

**RESPONSE: Excellent suggestions. We now present the requested experiments. The DNA content data are in Figure 3C, and the phospho-Rps6 data in the new Figure 4D are discussed in the text. Briefly, exogenous valine, and to a lesser extent leucine, suppressed the G1 accumulation, but not to wild type levels. Exogenous valine also substantially increased TORC1 activity (>10-fold).**

-It seems important to show that Rps6-P will decline in M phase and increase during a second cell cycle to establish that TORC1 activity actually fluctuates in the cell cycle instead of just by reduced by the manipulations involved in collecting young G1 cells by elutriation.

**RESPONSE: The second cycle comment is not pertinent to our elutriation setup. The two-cycle approach should be used in arrest-and-release synchronizations to minimize arrest-related artifacts when cells continue to grow in size. This is why we used elutriation in the first place, as described in the text, to avoid such artifacts. In elutriations it is the first cycle, exclusively of daughter cells, that can be meaningfully scored. After that, the cells lose synchrony very fast because you have mothers (which grow in size very little) and daughters (which need to double in size until mitosis). Hence, the second cycle will be meaningless and impossible to interpret.**

Reviewer #2 (Significance (Required)):

General Assessment:

Strengths: Evidence for BCAA biosynthesis from glucose in the G1 phase of the cell cycle, and evidence obtained from analyzing the bat1 mutant that BCAA synthesis underlies activation of TORC1 early in the cell cycle in a manner required to achieve the critical cell size necessary for G1 to S transition.

Weaknesses: Lack of evidence for Val biosynthesis in G1 despite evidence that Val limitation is more crucial than Leu limitation in the bat1 mutant; lack of confirmation that Val limitation underlies the delayed G1-S transition and reduced TORC1 signaling in the bat1 mutant; and lack of compelling evidence that TORC1 activity is periodic in WT cells.

Advance: This would be the first evidence that TORC1 activity varies through the cell cycle in a manner controlled by synthesis of BCAAs

Audience: This advance would be of great interest to a wide range of workers studying how the cell cycle is regulated and the role of TORC1 in controlling cell growth and division in normal cells and in human disease.

My expertise: Mechanisms of metabolic regulation of gene expression at the transcriptional and translational levels in budding yeast

**Referees cross-commenting**

Ref. #1's major comment 1 echoes my request for clarification about whether Leu, and not just Val, is limiting growth in the bat1 mutant, and also the need to determine which BCAA supplement to bat1 cells will restore TORC1 activity (which was also requested by Ref. #3).

I agree with this reviewer's request to provide evidence that Leu levels actually increase during G1 progression (comment #2). I also think the suggested experiments in Comment #3 are reasonable for their potential to provide stronger evidence that Leu production in the G1 phase of wild-type cells activates TORC1, as currently the argument is based on the finding of low TORC1 activation in bat1 cells (that seem to be limiting for Val vs. Leu). Comment #4 echoes similar requests made by both me and Ref. #3. Ref. #3's major comments 1 and 3 mirror two of my major comments. I wasn't convinced of the need to monitor Sch9 versus Rps6 phosphorylation as a read-out of TORC1 activity-does being a direct substrate truly matter? Regarding comment 5, I wasn't convinced of the need to include Rap-sensitive or -resistant control strains for the analysis in Fig. 5A. And regarding comment 4, while it would be interesting to examine if TORC1 regulates BCAA synthesis during cell cycle progression, this seems to be outside the scope of a demonstration that BCAA synthesis stimulates TORC1.

Thus, it seems we all agree on certain experiments that need to be carried out, and Ref. #1 has rightly proposed a few others with the potential to strengthen the evidence that Leu production during G1 phase mediates cyclic activation of TORC1

Reviewer #3 (Evidence, reproducibility and clarity (Required)):

In this manuscript, Blank and colleagues measure the synthesis of various metabolites from glucose during cell cycle progression and observe an increased synthesis of branched-chain amino acids (BCAA) from the early G1 to late G1 phase. Interestingly, they also found a gradual increase in TORC1 activity from the early G1 to the S phase which is proposed to be dependent on BCAA synthesis.

Major comments:

1. The authors show that TORC1 activity increases from the early G1 to the S phase. TORC1 activity is sensitive to short-term starvations caused during changing media or centrifugations. Hence, the concern arises regarding the increased pattern of TORC1 activity during the cell cycle. Is it really a biological phenomenon or a cellular adaptation to experimental conditions? Can authors provide more support for this observation? Can authors monitor the cell cycle for the two cell cycles to confirm that TORC1 activity shows a wavy pattern?

**RESPONSE: The same point was also made by Reviewer #2. As we noted in our response above, “The second cycle comment is not pertinent to our elutriation setup. The two-cycle approach should be used in arrest-and-release synchronizations to minimize arrest-related artifacts when cells continue to grow in size. This is why we used elutriation in the first place, as described in the text, to avoid such artifacts. In elutriations it is the first cycle, exclusively of daughter cells, that can be meaningfully scored. After that, the cells lose synchrony very fast because you have mothers (which grow in size very little) and daughters (which need to double in size until mitosis). Hence, the second cycle will be meaningless and impossible to interpret.”**

2. The authors use Rps6 phosphorylation as a read-out of TORC1 activity, which is not a direct substrate of TORC1. Analysis of the direct substrates of TORC1, such as phosphorylation of Sch9 will solidify the author's claim.

**RESPONSE: The reviewers discussed this point (see their comments above). We agree with the opinion that Rps6 phosphorylation accurately reports on TORC1 activity (also used in the fly experiments we now cite, as requested by Reviewer 1). For all our experiments' objectives and conclusions, it doesn't matter if the phosphorylation of Rps6 lies more downstream than Sch9 phosphorylation.**

3. Authors show that Bat1 lacking strain have reduced TORC1 activity. Can authors restimulate these cells with Leucin, Valine, and Isoleucine individually or in combination to identify the critical amino acid for the TORC1 activity?

**RESPONSE: Yes, that is an excellent suggestion. We show the experiment in Figure 4D (see previous response). Valine showed pronounced activation (>10-fold).**

4. The authors claim that increased BCAA synthesis is necessary for TORC1 activation. Since TORC1 is shown to be upstream of amino acid biosynthesis pathways, it will be interesting to check if TORC1 per se regulates BCAA synthesis during cell cycle progression. The authors could inhibit TORC1 by rapamycin treatment and monitor if the BCAA synthesis still shows cell cycle-dependent modulation.

**RESPONSE: The reviewers also discussed this point (see their comments above). We agree with the view that it is a very substantial undertaking, well beyond the scope of this work.**

5. In Figure 5A, the use of any rapamycin-sensitive and rapamycin-resistant strains as controls will strengthen their claim of TORC1 inhibition being epistatic to Bat1 deletion, since the rapamycin in minimal media might be less effective.

**RESPONSE: Again, the reviewers also discussed this point (see their comments above). We agree that it will not add much to the conclusions in the context of all the data we show and the existing literature.**

Minor comments:

1. The data of metabolic labeling, especially various species M1, M2, M3, etc., of an individual metabolite is difficult to understand for the general readers. Hence, a schematic explaining various species might be helpful.

**RESPONSE: We added a new Figure (EV1) delineating the carbons from glucose to valine and leucine.**

2. Please describe the elutriation approach in more detail with media conditions and buffer conditions to understand the overall experimental setup.

**RESPONSE: We now added this information (see the second section of the Materials and Methods).**

Reviewer #3 (Significance (Required)):

Significance:

Overall, this study presents an interesting observation to the researchers working in TORC1 and cell cycle regulation.
