## Supplementary material for "Branched chain amino acid synthesis is coupled to TOR activation early in the cell cycle in yeast": Marked-up file with text changes

Figure EV3:	Alpha-keto acid supplementation does not suppress the growth defect of *bat1Δ* mutants.

Figure EV4:	Amino acid composition of *BAT1^+^* and *bat1Δ* strains from PTH-based analysis.

Figure EV5:	Volcano plot of steady state abundances of primary metabolites and biogenic amines.

Figure EV6:	Cells lacking Bat1 are smaller, increase in size slower, and are delayed in the G1 phase of the cell cycle.

File S1:	Source data for the graphs in all Figures.

File S2:	Metabolite measurements from the GC-MS measurements during the cell cycle, from the isotope tracing experiments.

File S3:	Steady-state primary metabolite measurements in *BAT1^+^* vs. *bat1Δ* cells.

File S4:	Steady-state biogenic amine measurements in *BAT1^+^* vs. *bat1Δ* cells.

File S5:	Source immunoblots for Figures 4B,C.

File S6:	Source immunoblots for Figure 4D.

File S7:	Source immunoblots for Figures 5A,B.

File S8:	Additional source immunoblots for Figure 5B.

File S9:	Reagent Table

**ACKNOWLEDGEMENTS**

We thank Maria Cardenas-Corona (NIH) for reagents. This work was supported by the National Institutes of Health (NIH, grant R01 GM123139) and by the German Research Foundation (DFG, Project-ID 34509606—TRR 51).

**AUTHOR CONTRIBUTIONS**

M.P. and K.H. conceived and designed the study. C.R. and K.S-H. performed and analyzed the isotope tracing experiments. S.E.H. performed the cell cycle analyses shown in Figure 3C. H.B. and M.P. performed all other experiments. All authors evaluated and discussed the data. M.P., H.B., and C.R. wrote the first draft of the manuscript, which was then edited by all authors.

**CONFLICTS OF INTEREST**

The authors have no conflicts of interest to declare that could be perceived to influence the presentation or interpretation of the data.

**FIGURES**

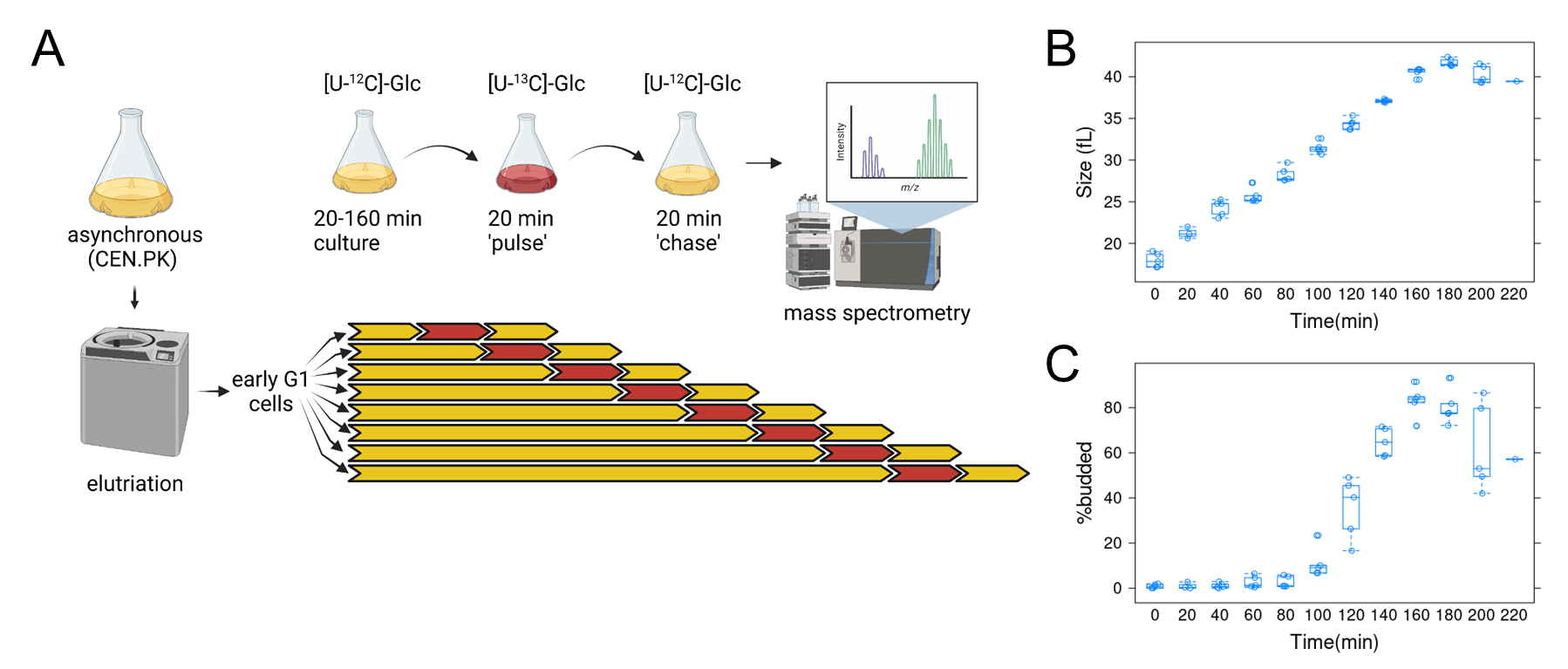

**FIGURE 1.** **Overview of the approach to obtain samples for metabolic flux analysis in highly synchronous, dividing, budding yeast cells.** A, For each experiment, early G1 daughter cells of a prototrophic strain (CEN.PK; see Materials and Methods) were obtained by centrifugal elutriation. The elutriated culture was split into eight aliquots and cultured for a varying amount of time, from 20 to 160 min, in minimal, [U-^12^C]-glucose medium. They were then transferred to a medium with [U-^13^C]-glucose for 20 min (pulse) and then incubated for another 20 min in [U-^12^C]-glucose medium (chase). Metabolite extracts from these cells were analyzed by mass spectrometry. Five such independent experiments were performed. The figure was generated with Biorender.com. B, Boxplots showing the cell size (y-axis) over time (x-axis) of all the samples as they progressed in the cell cycle. C, Boxplots showing the percentage of budded cells (y-axis) from the same samples shown in B. The boxplot graphs were generated with R language functions. The values used to draw the plots are in File S1/Sheet1.

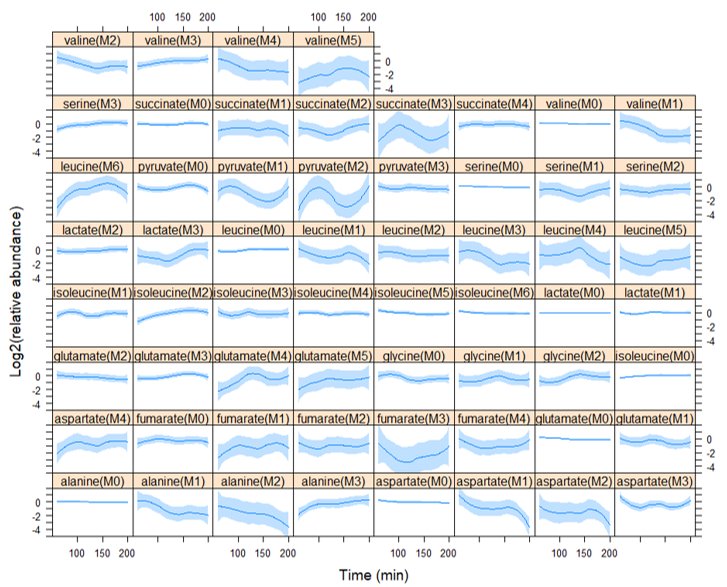

**FIGURE 2.** **Relative abundance of intracellular metabolite isotopomers in the cell cycle.** Each plot shows the relative mass isotopomer distribution (MID) values for metabolite species detected from cell extracts in each of the five independent experiments shown in Figure 1. Each value was divided by the average value of the entire time series (x-axis) for that species from the same experiment, and Log2-transformed (y-axis). Loess curves and the std errors at a 0.95 level are shown. The values used to generate the graphs are in File S1/Sheet2. The Log2(relative abundance) values of pyruvate(M1) and leucine(M6) showed the most significant changes (p<0.1, see File S2/Sheet2) in cells across the cell cycle.

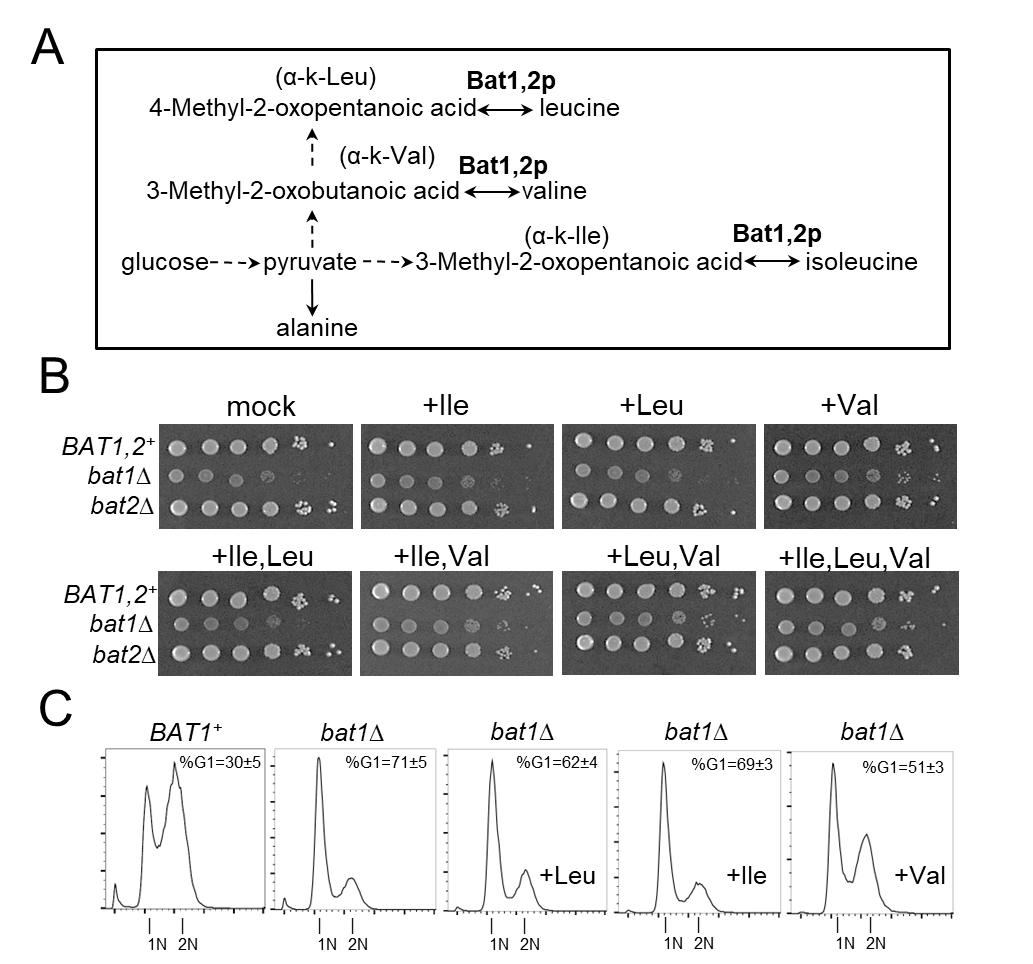

**FIGURE 3. BCAA supplementation suppresses the growth defect of *bat1Δ* mutants.** A, Diagram of the reactions leading to BCAAs from the corresponding alpha-keto (α-k) acids, catalyzed by Bat1,2p. A more detailed diagram leading to the of valine (M5) and leucine (M6) isotopomers is in Figure EV1. B, The indicated strains (all in the prototrophic CEN.PK background; see Materials and Methods) were spotted at 10-fold serial dilutions on solid Synthetic Minimal Medium (SMM) agar plates. Exogenous amino acids were added at 1mM final concentration, as indicated in each case. The plates were incubated at 30 °C and photographed after 3-days. C, DNA content profiles of *BAT1* and *bat1Δ* cells from asynchronous cultures, in SMM medium. Where indicated, exogenous amino acids were added at 1mM final concentration. On the x-axis of the histograms is fluorescence per cell, while on the y-axis is the cell number. Peaks corresponding to cells in G1 with unreplicated (1N) and cells in G2 and M phases with fully replicated (2N) DNA are indicated. The percentage of cells with G1 DNA content (%G1) from 3 independent measurements is shown in each case (mean and sd). The values used to generate the graphs are in File S1/Sheet3.

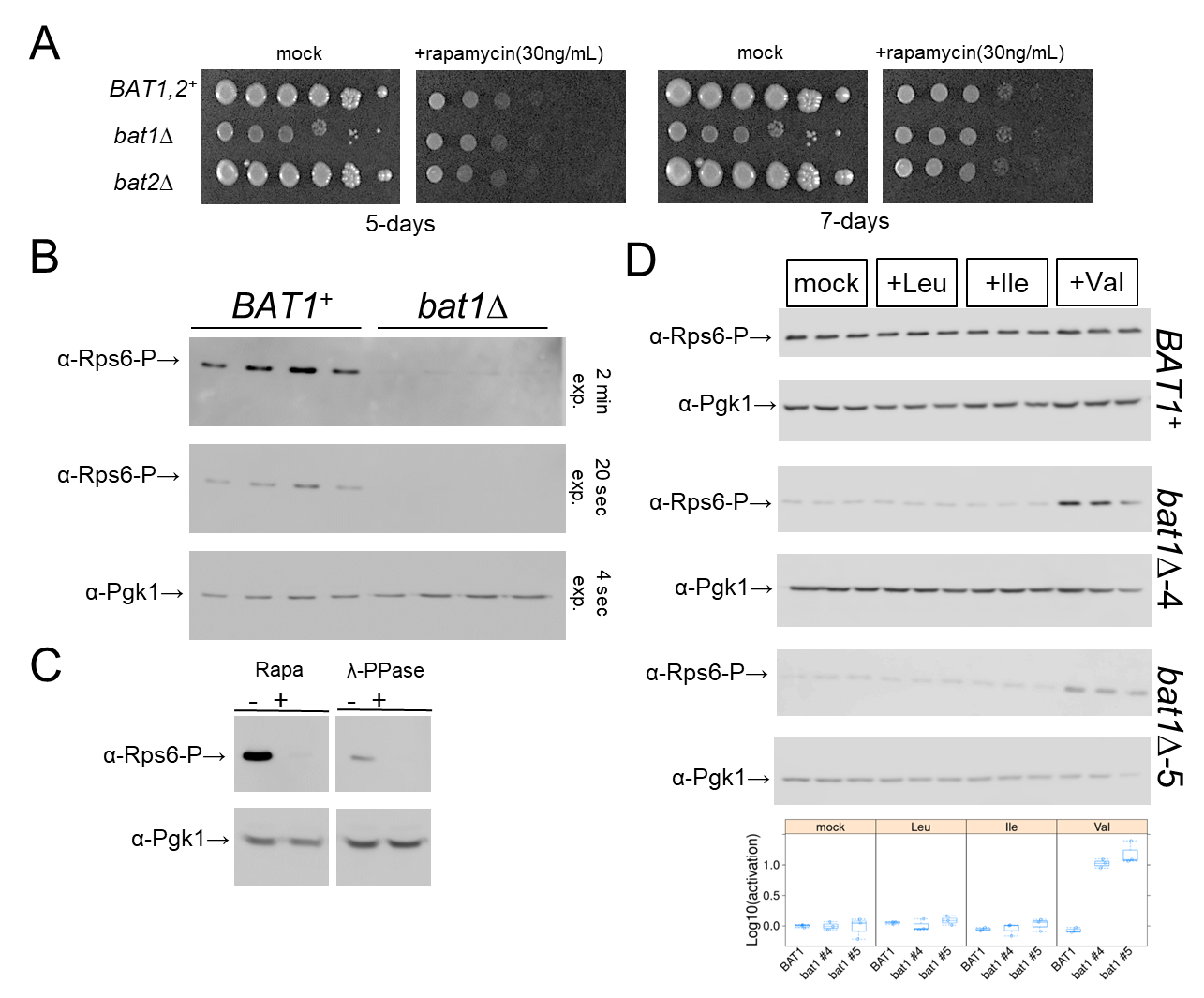

**FIGURE 4. Bat1 is functionally linked to TORC1 activation in early G1.** A, The indicated strains were spotted at 10-fold serial dilutions on solid Synthetic Minimal Medium (SMM) agar plates. Rapamycin was added at 30 ng/mL, as shown in each case. The plates were incubated at 30 °C and photographed after 5 and 7 days, as indicated. B, Immunoblots of total cell extracts from asynchronous *BAT1* and *bat1Δ* cells, from four independent experiments. The signal from Pgk1 (α-Pgk1) is shown on the blot at the bottom, and from phosphorylated Rps6 (α-Rps6-P) is on the blot above, indicated for two exposures (2 min, top; 20 sec, middle). C, On the left are immunoblots from wild-type (CEN.PK) cells treated (+), or not (-), with rapamycin (Rapa) at 200 ng/mL for 1 h before cell extract preparation. On the right are immunoblots of wild-type (CEN.PK) cell extracts. The extracts were treated (+), or not (-), with λ-phosphatase for 1 h (see Materials and Methods). The levels of phosphorylated Rps6 (α-Rps6-P) and Pgk1 (α-Pgk1) are shown in each case. The raw immunoblots for B & C are in File S5. D, Exogenous addition of valine leads to sustained activation of TORC1 and phosphorylation of Rps6 in cells lacking Bat1. Wild type (*BAT1^+^*) and *bat1∆* (two independent isolates, #4, and #5) strains were grown overnight in minimal (SMM) medium, diluted to 1E+06 cells/mL in fresh SMM media containing the indicated amino acid (at 1mM), and harvested when they reached 5E+06 cells/mL. The levels of phosphorylated Rps6 (α-Rps6-P) and Pgk1 (α-Pgk1) are shown in immunoblots from total cell extracts in each case. The relative levels of Rps6-P/Pgk1 are shown in each case at the bottom. The values used to generate the graphs are in File S1/Sheet4. The raw immunoblots for D are in File S6.

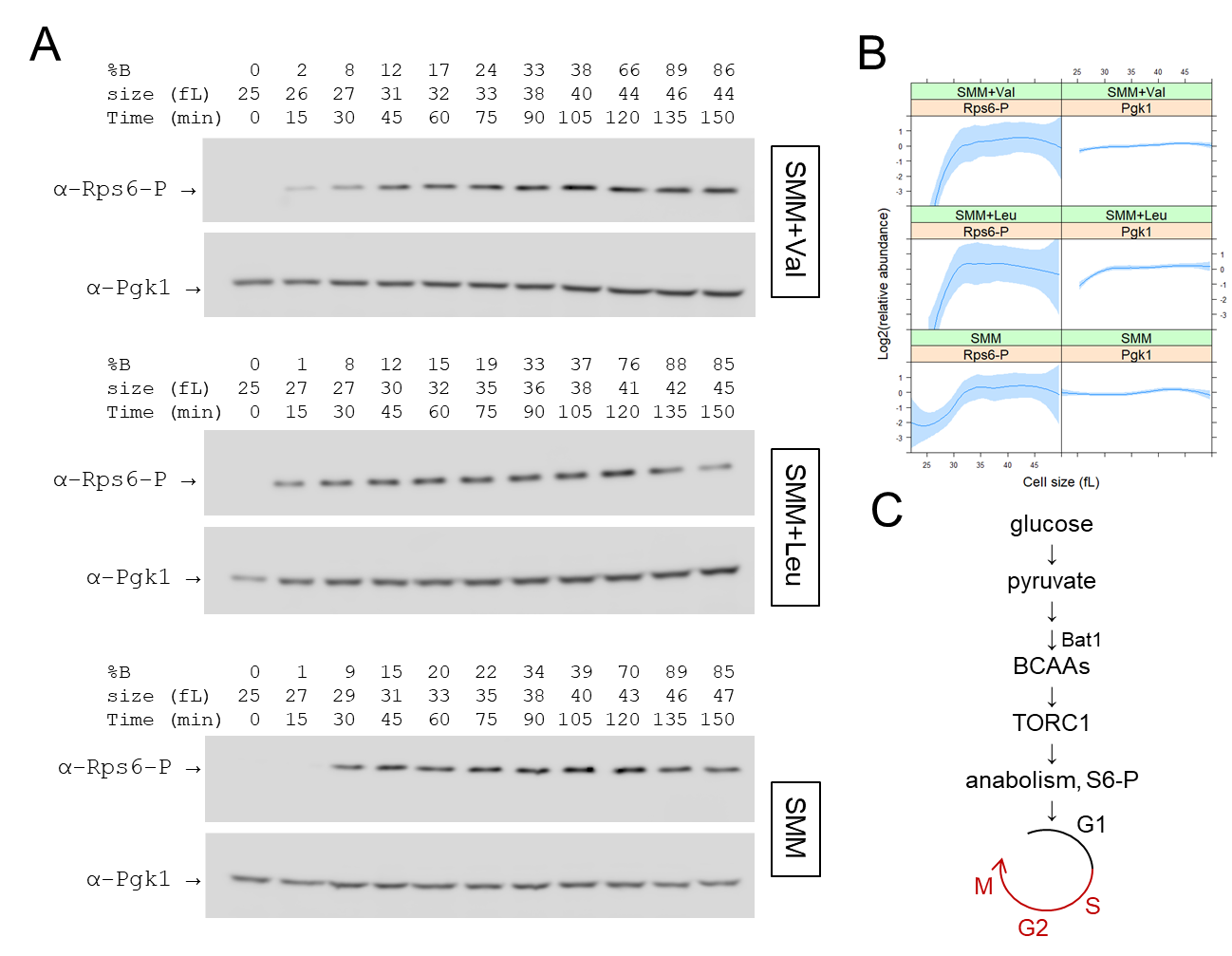

**FIGURE 5. TORC1 activity increases as cells progress in the cell cycle.** A, Immunoblots of total cell extracts from synchronous, elutriated wild-type (CEN.PK) cells in minimal (SMM) medium with exogenous Leu or Val added at 1mM immediately after elutriation. At the top, the percent of budded cells (%B), cell size (in fL), and time (in min) are indicated. The levels of phosphorylated Rps6 (α-Rps6-P) and Pgk1 (α-Pgk1) are shown in each case. B, Quantification of the levels of phosphorylated Rps6 and Pgk1 from independent experiments done as in A (see Files S7,8). The relative levels of each protein across the cell cycle is shown on the y-axis, as a function of cell size (x-axis). Loess curves and the std errors at a 0.95 level are shown. The values used to draw the plots are in File S1/Sheet5. C, Schematic of a possible model to explain our findings, linking BCAA synthesis to TORC1 activation early in the cell cycle.

**EXPANDED VIEW FIGURES**

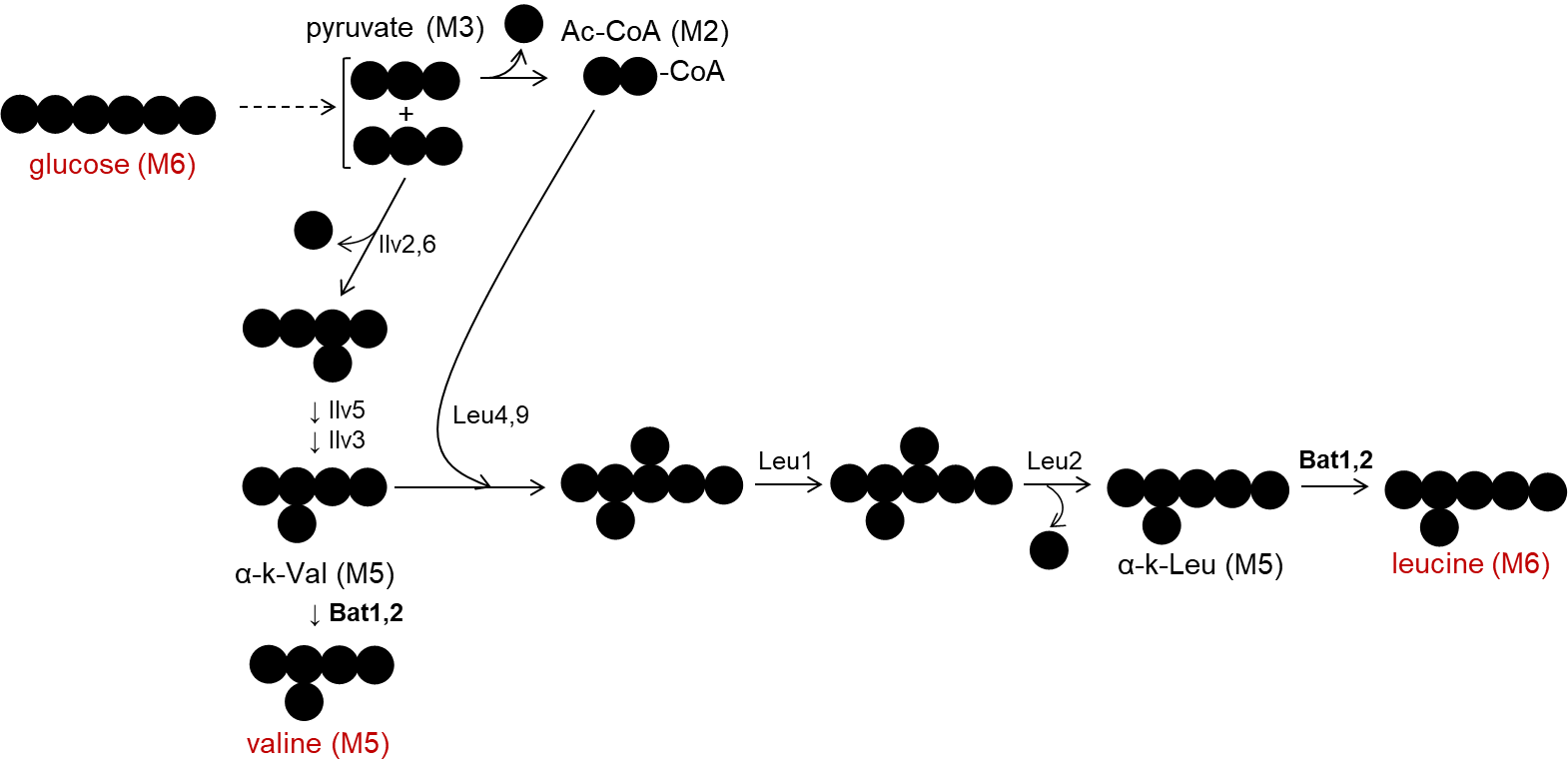

**FIGURE EV1. Diagram showing the carbon additions and eliminations from glucose (M6) to the synthesis of valine (M5) and leucine (M6) isotopomers.**

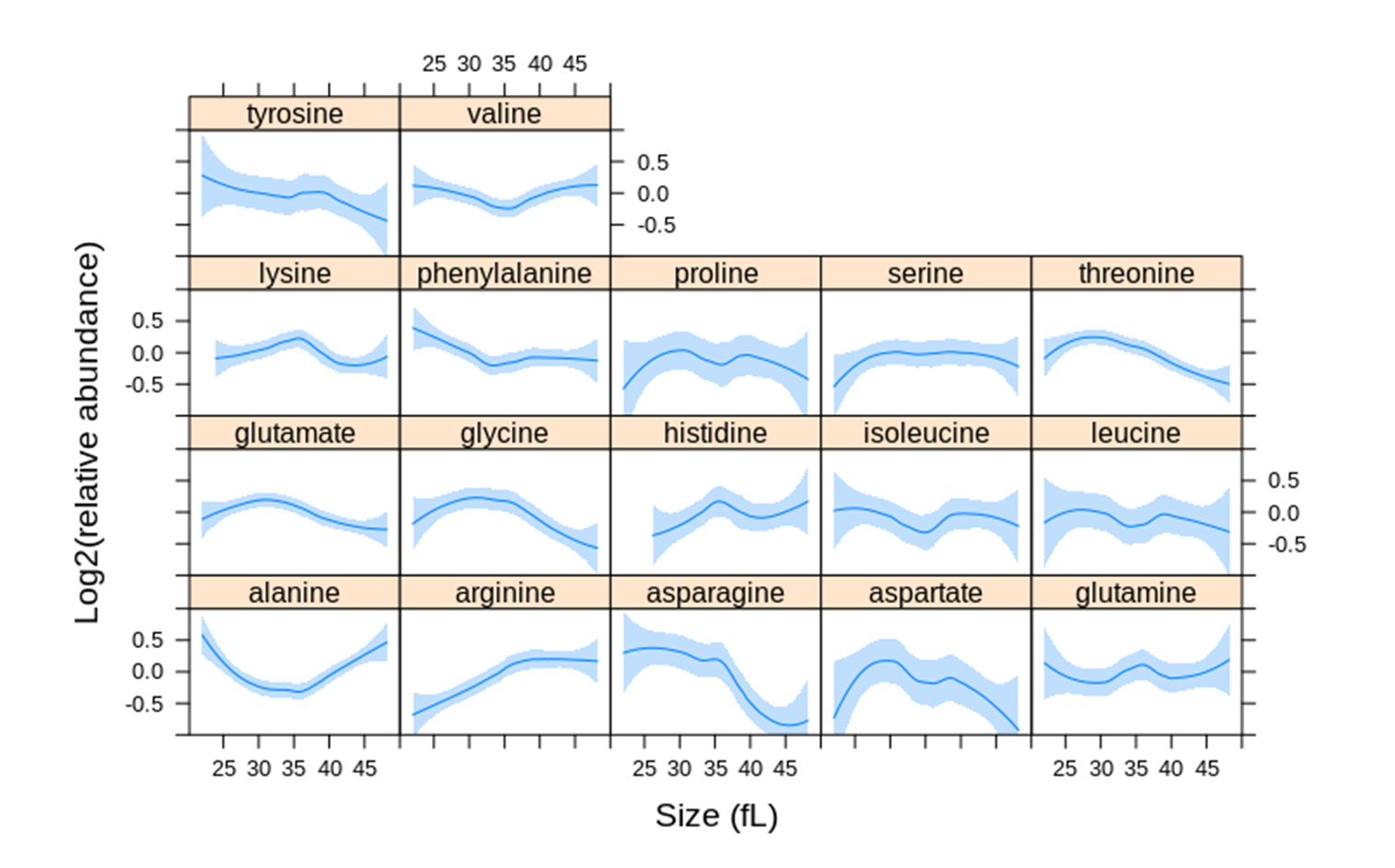

**FIGURE EV2. Steady-state amino acid levels in the cell cycle.** Intracellular amino acid levels were measured as described in Materials and Methods from synchronous, elutriated cultures in SMM media. The fraction of each amino acid is on the y-axis. The values used to draw the plots are in File S1/Sheet6.

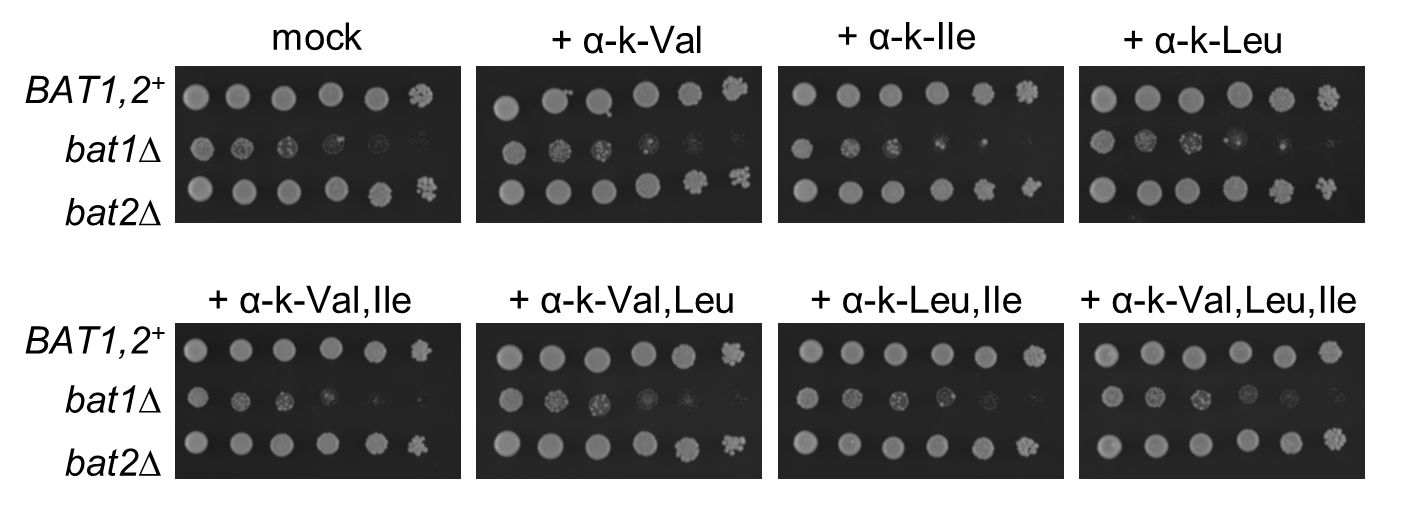

**FIGURE EV3. Alpha-keto acid supplementation does not suppress the growth defect of *bat1Δ* mutants.** The experiment was done as in Figure 3. Exogenous alpha-keto acids were added at 1mM final concentration, as indicated in each case.

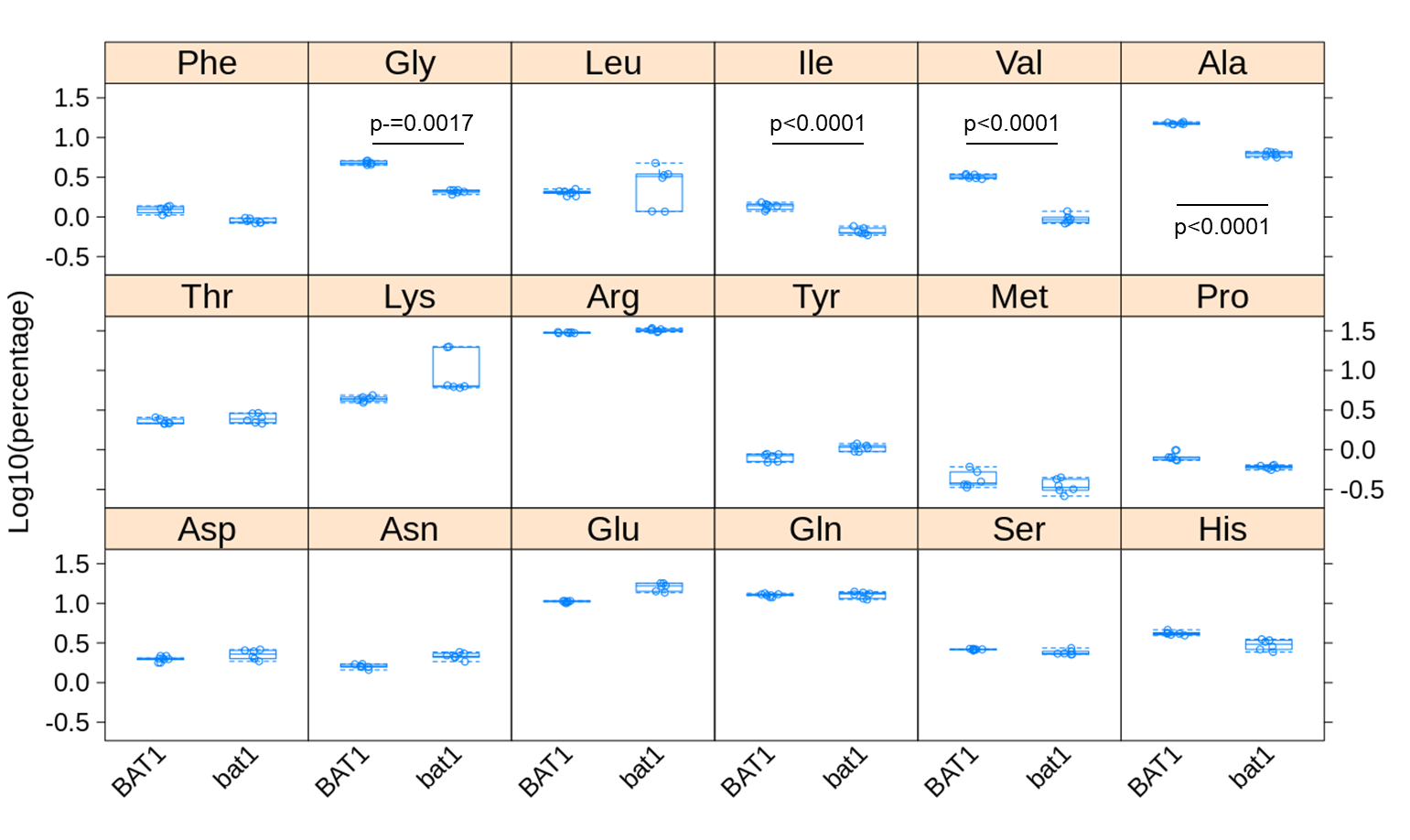

**FIGURE EV4. Amino acid levels in *BAT1^+^* and *bat1Δ* cells.** Intracellular amino acid levels were measured as described in Materials and Methods and Figure EV2, from exponentially growing cultures in SMM media. The fraction of each amino acid is on the y-axis. The four amino acids (Gly, Ile, Val, Ala) with significantly altered fractional abundance (>2-fold, p<0.05; from six independent samples in each case) are indicated. The values used to draw the plots are in File S1/Sheet7.

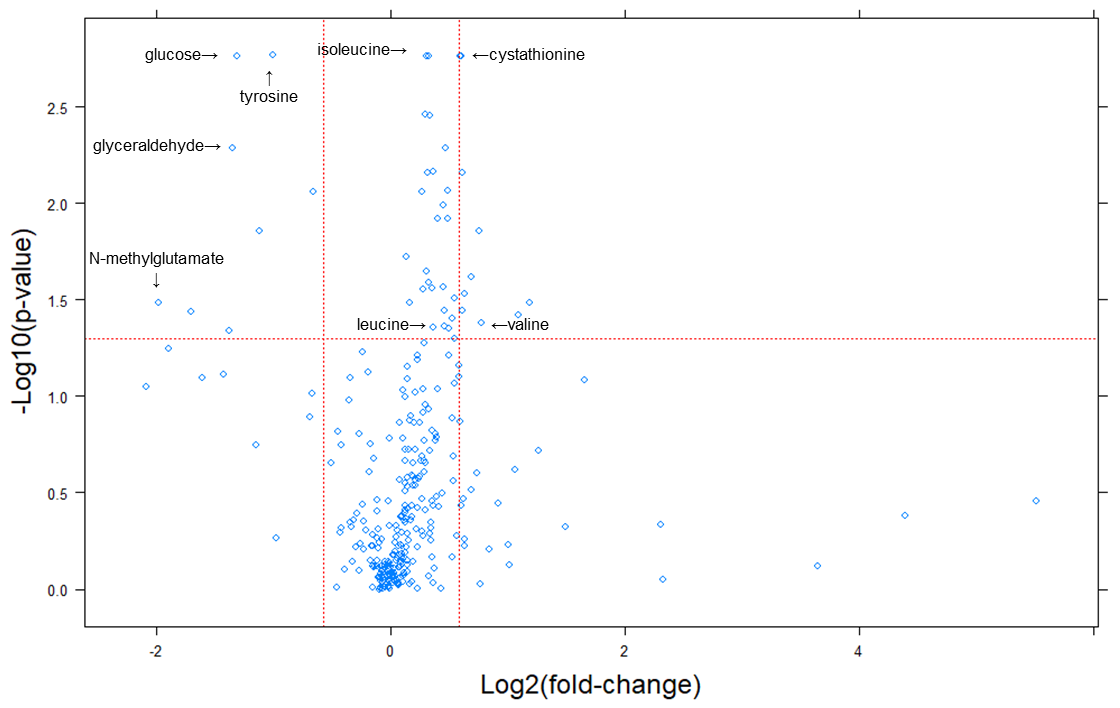

**FIGURE EV5. Changes in primary and biogenic amine metabolites in cells lacking Bat1.** Metabolites whose levels changed were identified from the magnitude of the difference (x-axis; Log2-fold change in *BAT1^+^* : *bat1Δ* cells) and statistical significance (y-axis), indicated by the red lines. The analytical and statistical approaches are described in Materials and Methods. The values used to generate the graphs are in File S1/Sheet8. The labels indicate selected metabolites.

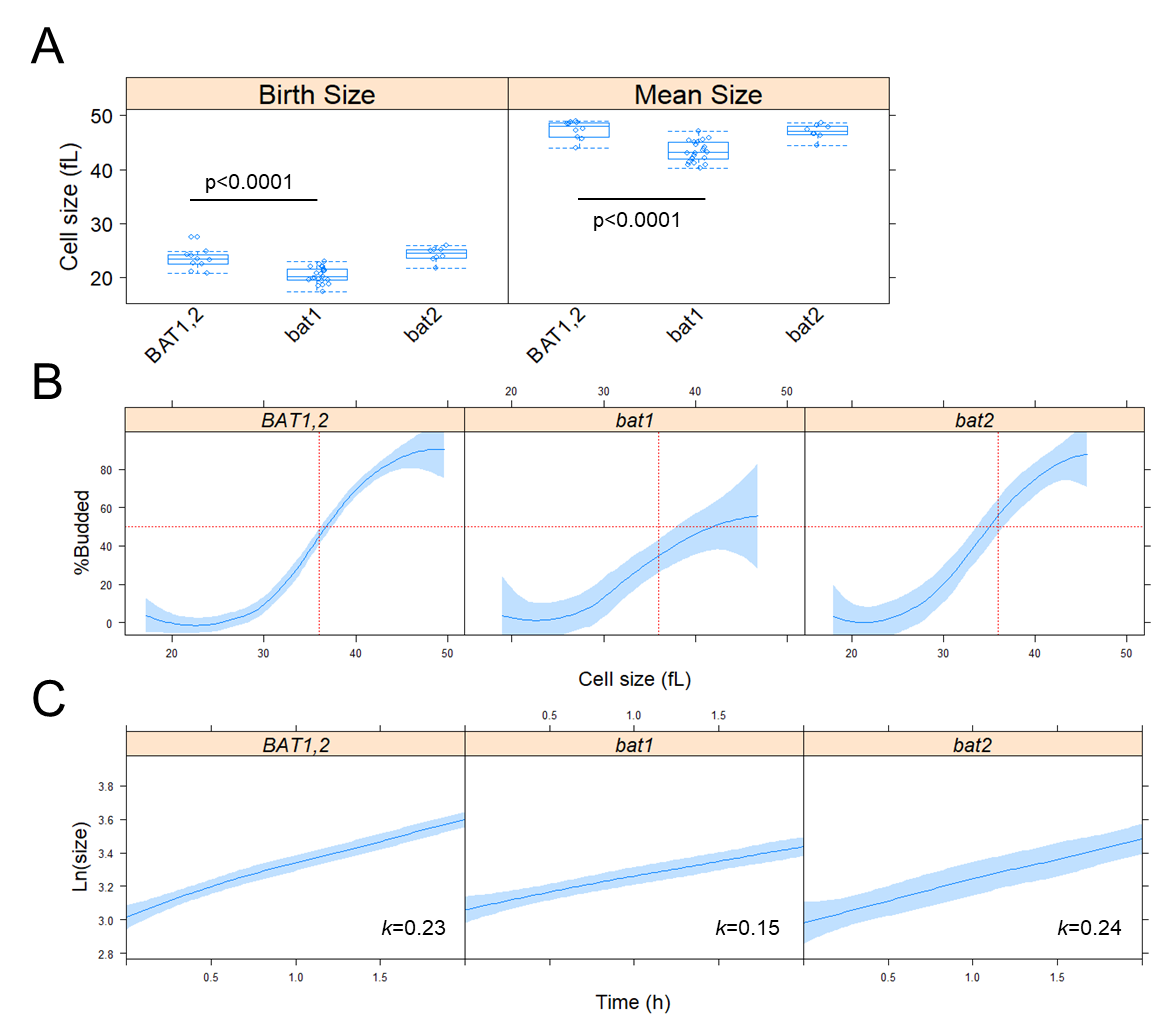

**FIGURE EV6.** **Cells lacking Bat1 are smaller, increase in size slower, and are delayed in the G1 phase of the cell cycle.** A. Boxplots showing the size of the cells (in fL). The measurements were taken from asynchronous cultures in synthetic minimal media (SMM) without amino acid supplementation. The values used to draw the plots are in File S1/Sheet9. B, Plots of the percentage of budded cells (y-axis) as a function of size (x-axis). The measurements were from daughter cells of the indicated strains, obtained by centrifugal elutriation, progressing in the cell cycle in SMM medium. Loess curves and the std errors at a 0.95 level are shown. C, From the same experiments as in B, the specific rate of the increase in size (*k*) was calculated, from the slope of the regression lines plotting the Ln-transformed cell size values (y-axis) against time (x-axis). The values used to draw the plots in B and C are in File S1/Sheet10.
